# Design of a highly thermotolerant, immunogenic SARS-CoV-2 spike fragment immunogen

**DOI:** 10.1101/2020.08.15.252437

**Authors:** Sameer Kumar Malladi, Randhir Singh, Suman Pandey, Savitha Gayathri, Kawkab Kanjo, Shahbaz Ahmed, Mohammad Suhail Khan, Parismita Kalita, Nidhi Girish, Aditya Upadhyaya, Poorvi Reddy, Ishika Pramanick, Munmun Bhasin, Shailendra Mani, Sankar Bhattacharyya, Jeswin Joseph, Karthika Thankamani, V. Stalin Raj, Somnath Dutta, Ramandeep Singh, Gautham Nadig, Raghavan Varadarajan

**Affiliations:** Molecular Biophysics Unit (MBU), Indian Institute of Science, Bengaluru, India; Mynvax Private Limited, ES12, Entrepreneurship centre, SID, Indian Institute of Science, Bengaluru, India; Translational Health Science and Technology Institute, NCR Biotech Science Cluster, 3rd Milestone, Gurugram-Faridabad Expressway, Faridabad-121001; Virology Scientific Research (VSR) Laboratory, School of Biology, Indian Institute of Science Education and Research Thiruvananthapuram (IISER TVM), Kerala, India; Jawaharlal Nehru Centre for Advanced Scientific Research, Jakkur, Bengaluru, India

**Keywords:** glycosylation, microbial, Pichia, thermostable, ACE2

## Abstract

Virtually all SARS-CoV-2 vaccines currently in clinical testing are stored in a refrigerated or frozen state prior to use. This is a major impediment to deployment in resource-poor settings. Several use viral vectors or mRNA. In contrast to protein subunit vaccines, there is limited manufacturing expertise for these novel, nucleic acid based modalities, especially in the developing world. Neutralizing antibodies, the clearest known correlate of protection against SARS-CoV-2, are primarily directed against the Receptor Binding Domain (RBD) of the viral spike protein. We describe a monomeric, glycan engineered RBD protein fragment that is expressed at a purified yield of 214mg/L in unoptimized, mammalian cell culture and in contrast to a stabilized spike ectodomain, is tolerant of exposure to temperatures as high as 100°C when lyophilized, upto 70°C in solution and stable for over four weeks at 37°C. In prime:boost guinea pig immunizations, when formulated with the MF59 like adjuvant AddaVax™, the RBD derivative elicited neutralizing antibodies with an endpoint geometric mean titer of ~415 against replicative virus, comparing favourably with several vaccine formulations currently in the clinic. These features of high yield, extreme thermotolerance and satisfactory immunogenicity suggest that such RBD subunit vaccine formulations hold great promise to combat COVID-19.

## Introduction

SARS-CoV-2 is the etiological agent of the on-going COVID-19 pandemic (1, 2). As on 4^th^ October, 2020 there are ~34.7 million infections and ~1.0 million deaths worldwide (3). The major surface protein of SARS-CoV-2 is the spike glycoprotein. Like several other viral surface glycoproteins, it is a homotrimer, with each protomer consisting of two subunits S1 and S2. The S1 subunit consists of an N-terminal domain (NTD), linker and receptor binding domain (RBD), and two small subdomains SD1 and SD2 (4–6) (Figure 1A–1D). The RBD domain of the spike glycoprotein binds to the cell surface receptor ACE2, followed by endocytosis or fusion mediated via the fusion peptide located on the S2 subunit (7). Most of the neutralizing antibody responses are targeted to the RBD (8–14), though very recently, neutralizing antibodies against the NTD have also been identified (15). It is thus unclear whether the full length spike or the RBD is a better immunogen.

**Figure 1:**
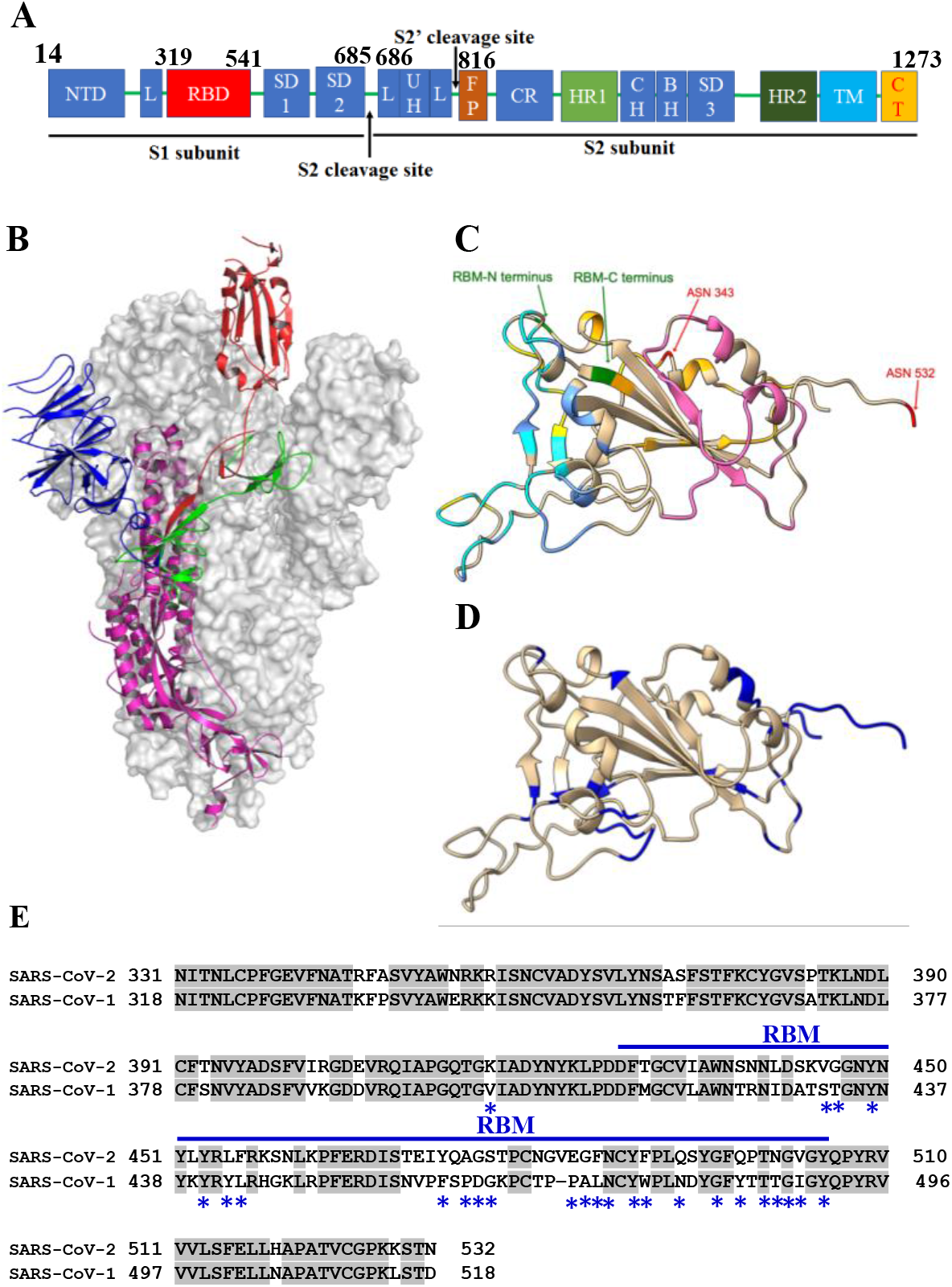
S-protein domain organization, structure of Spike and receptor binding domain of SARS-CoV-2. **A)** Linear map of the S protein spike with the following domains: NTD, N-terminal domain; L, linker region; RBD, receptor-binding domain; SD, subdomain; UH, upstream helix; FP, fusion peptide; CR, connecting region; HR, heptad repeat; CH, central helix; BH, β-hairpin; TM, transmembrane region/domain; CT, cytoplasmic tail. **B)** Spike ectodomain trimer highlighting protomer with RBD in the ‘up’ conformation, NTD in dark blue, RBD in brick red, SD1 and SD2 in green and S2 subunit in magenta (PDB: 6VSB) **C)** Epitopes for known RBD directed neutralizing antibodies. The N and C termini of the receptor-binding motif (RBM) are labeled and in green. Residues at the binding interfaces with hACE2 are in cyan. The B38 epitope has considerable overlap with the hAce2 interface, non-overlapping residues are in light blue. Epitopes for S309, P2B-2F6 are in orange and yellow. Epitope for CR3022 is in pink, this overlaps substantially with the potent neutralizing antibody H014. The conserved N-glycosylation site at 343 and the engineered, immune masking glycosylation site at 532 are shown in red. **D)** Exposed residues with solvent accessible surface area that are not part of any neutralizing epitope identified so far are shown in dark blue. The largest such stretch is at the C-terminus where the engineered glycosylation site is placed. **E)** Sequence alignment of SARS-CoV-1 (residues: 318-518) and SARS-CoV-2 (residues 331-532), the blue line indicates the receptor binding motif (RBM), the grey highlight indicate residues conserved in both SARS-CoV-1 and SARS-CoV-2 and the blue asterisks indicate the ACE2 binding residues. (PDB: 6M0J)

Over 150 vaccine candidates are under development globally (16). Some vaccine candidates that have entered rapidly into clinical phase testing include mRNA vaccine candidates by Moderna (mRNA-1273), BioNTech (BNT162b1)(17), and CureVac (CVnCoV), a Chimpanzee Adenovirus vector vaccine by University of Oxford and AstraZeneca (ChAdOx1-S) (18), a non-replicating adenovirus type-5 (Ad5) vaccine by Cansino (Ad5-nCoV) (19), a DNA vaccine by Inovio (INO-4800) (20), inactivated virus vaccines by Sinovac (PiCoVacc) (21) and Bharat Biotech (COVAXIN), a native like trimeric subunit spike protein vaccine by Clover Biopharmaceuticals /GSK/Dynavax (SCB-2019), and a full length recombinant glycoprotein nanoparticle vaccine by Novavax (NVX-CoV2373) (16, 22). The majority of the above employ full length spike or the corresponding ectodomain as the antigen. While there is some encouraging pre-clinical and Phase 1 clinical data, there is no precedent for use of mRNA or viral vectors, which are the farthest along in clinical development, in mass human vaccinations. In addition, with inactivated or attenuated virus, there are obvious safety issues that need careful attention. There are few studies that compare the relative immunogenicity of multiple vaccine candidates expressed in multiple platforms (23). Herein, we report a mammalian cell expressed, glycan engineered, RBD based subunit vaccine candidate (mRBD) formulated with an MF59 equivalent adjuvant. In contrast to an equivalent *Pichia pastoris* expressed RBD protein formulation, mRBD elicits titers of neutralizing antibodies in guinea pigs well above the levels required for protection in non-human primate challenge studies. mRBD expresses at eight-fold higher levels and is substantially more tolerant to thermal stresses than a stabilized spike ectodomain without compromising immunogenicity, and can be stored for over four weeks at 37°C. These data suggest that it is a promising candidate for further clinical development.

## Results

### Design of a recombinant RBD subunit vaccine

The RBD of the spike protein is the major target of neutralizing antibodies (8, 10–14, 24). SARS-CoV-2 is 79.6% identical to SARS-CoV-1 sequences (25). The spike protein of SARS-CoV-2 is 80% identical to its homolog from SARS-CoV-1. The RBD of SARS-CoV-2 shares 74% amino acid sequence identity with the RBD of SARS-CoV-1. We hypothesized that a receptor binding domain subunit derivative that lacks flexible termini as well as unpaired cysteines, and retains the ACE2 receptor binding site, located within the receptor binding motif (RBM, comprising residues 438-505, Figure 1E) as well as the cryptic epitope recognized by the neutralizing antibody CR3022 would be a good immunogen. We selected the RBD residues based on SWISS Model structure-based modelling of SARS-CoV-2 sequence, prior to availability of any SARS-CoV-2 spike and RBD-ACE2 complex structures. The modelled structure closely resembles the recently determined experimental structures determined by X-ray crystallography or Cryo-EM (RMSD: 1.2Å, with X-ray structure PDB: 6M0J) (5, 6, 26). In the X-ray structure, residues after 526 are disordered. Two RBD sequences were shortlisted consisting of residues 331-532 and 332-532 with retention (m331RBD) or deletion (mRBD/pRBD) of the native glycan at N331 for expression in mammalian and *P. pastoris* expression systems respectively. The constructs for mammalian expression are designated as m331RBD and mRBD, and for *Pichia* expression, pRBD respectively. In the past few months, several potent neutralizing antibodies directed against the RBD have been isolated and it currently appears that virtually the entire exposed surface of the RBD is targeted by neutralizing antibodies, with the exception of the C-terminal region distal from the RBM. We have introduced a glycosylation site at N532 in all the above RBD constructs to mask this region of the surface (Figure 1C, 1D).

### RBD (332-532) is more highly expressed and thermotolerant than a stabilized spike ectodomain

Mammalian cell expressed m331RBD and mRBD were purified by single step Ni-metal affinity chromatography from transiently transfected Expi293F culture supernatants. The proteins were confirmed to be predominantly monomeric by SEC (Figure 2A). Proteins from both the constructs were pure and were expressed at yields of ~68 ± 10 mg/L and ~214 ± 9 mg/L for m331RBD and mRBD respectively. Removal of the N-terminal glycan in m331RBD by introducing the T333H mutation resulted in substantially increased expression, similar to that of mRBD, confirming that the presence of the N-terminal glycan is responsible for reduced yield, as has been observed previously for SARS-CoV-1 RBD (27). All proteins were monomeric. In SEC, m331RBD which has an additional glycan, elutes before mRBD (Figure 2A). Given the higher yield of mRBD, most subsequent studies were carried out with this RBD derivative. nanoDSF thermal melt studies demonstrated that removal of the N-terminal glycan did not affect protein stability (Figure 2B). mRBD bound ACE2-hFc with a K_D_ of about ~14.2 nM (Figure 2C) and the neutralizing antibody CR3022 with a K_D_ of 16 nM, confirming that the molecule is properly folded (Figure 2D). mRBD is digested by trypsin with approximate half-lives of 20 and 60 minutes at 37 and 4 °C (Figure 2E) respectively. The digestion kinetics is unaffected by storage for over a week at 4 °C.

**Figure 2:**
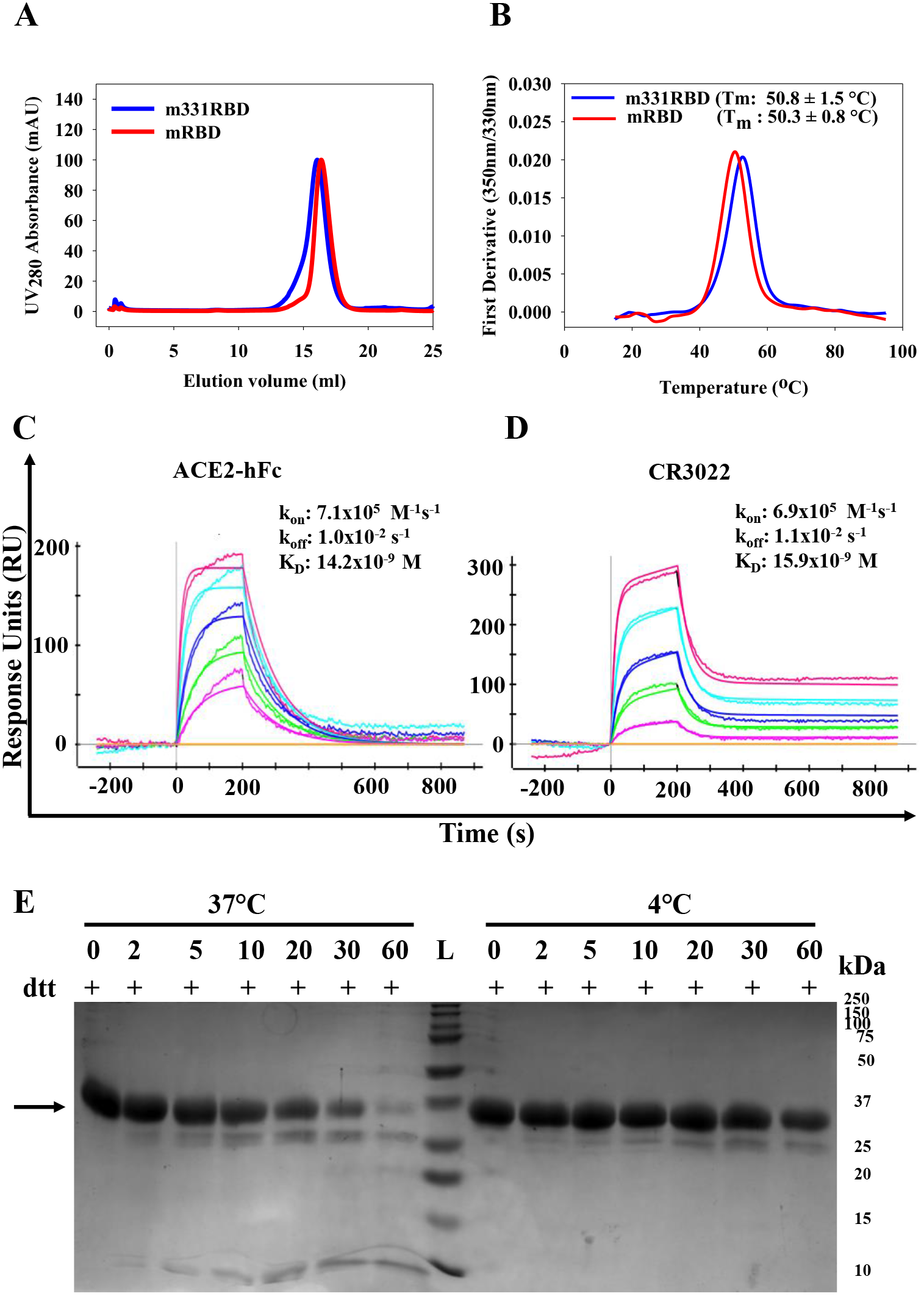
Characterization of mammalian cell expressed RBD. **A)** Size exclusion chromatography profile of m331RBD, mRBD immunogens with predominantly monomeric peak at ~16.0 and ~16.3mL respectively on S200 10/300GL column calibrated with Biorad gel filtration marker (Cat.No. 1511901) run at flowrate of 0.5mL/min with PBS (pH 7.4) as mobile phase **B)** nanoDSF equilibrium thermal unfolding of m331RBD and mRBD. **C)** SPR binding sensorgrams to ACE2 receptor. The concentrations of mRBD used as analytes are 100 nM, 50 nM, 25 nM, 12.5 nM, 6.25 nM **D)** SPR binding sensorgrams of mRBD with the neutralizing antibody CR3022. mRBD analyte concentrations are 50 nM, 25 nM, 12.5 nM, 6.2 nM, and 3.1 nM. **E)** Limited proteolysis of purified mRBD protein by TPCK treated trypsin (RBD:TPCK Trypsin=50:1) at 4°C and 37°C.

A construct with identical amino acid sequence to mRBD (pRBD) was expressed and purified from *P. pastoris* strain *X-33* from a stably integrated gene cassette at a yield of ~50 mg/L in shake flasks. The *Pichia* protein is more heterogeneous, extensively glycosylated and elutes at higher molecular weight than mRBD in both SDS-PAGE and SEC (Supplementary Figure 1A, 1F). The thermal stability of the *Pichia* purified immunogen pRBD (T_m_: 49.2 °C) is similar to mammalian cell expressed versions (Supplementary Figure 1B). The protein bound with comparable affinity to ACE2-hFc and CR3022 with K_D_’s of approximately 23 nM and 30 nM respectively, similar but slightly higher than corresponding values for mRBD (Supplementary Figure 1D, 1E). *Pichia* expressed RBD was similarly stable to thermal stress and proteolysis (Supplementary Figure 1C, 1F, 1G). We also attempted to express the protein in *E. coli*. The protein expressed well but was targeted to inclusion bodies. Despite multiple attempts employing a variety of refolding strategies, we were unable to obtain significant quantities of properly refolded protein, competent to bind ACE2 from *E. coli*.

The spike ectodomain and full length spike formulations are important SARS-CoV-2 vaccine candidates (5, 6, 22) and it is therefore important to compare mRBD with these. We purified the Spike-2P stabilized ectodomain (Spike containing mutations K968P and V969P) protein from Expi293F cells by single step nickel chelate affinity chromatography followed by tag removal with a purified yield of ~25 mg/L culture (5). The purified protein was observed to be trimeric on SEC and bound tightly to ACE2-hFc with little dissociation (Figure 3A, 3B). Negative-stain EM confirmed that Spike-2P purified by us adopts a native like elongated trimeric structure (Figure 3C) consistent with available structures determined by Cryo-EM (5, 6, 15). Spike-2P was rapidly digested by trypsin with approximate half-lives of 10 and 30 minutes at 37 and 4 °C respectively (Figure 3D–E) yielding multiple RBD containing fragments.

**Figure 3:**
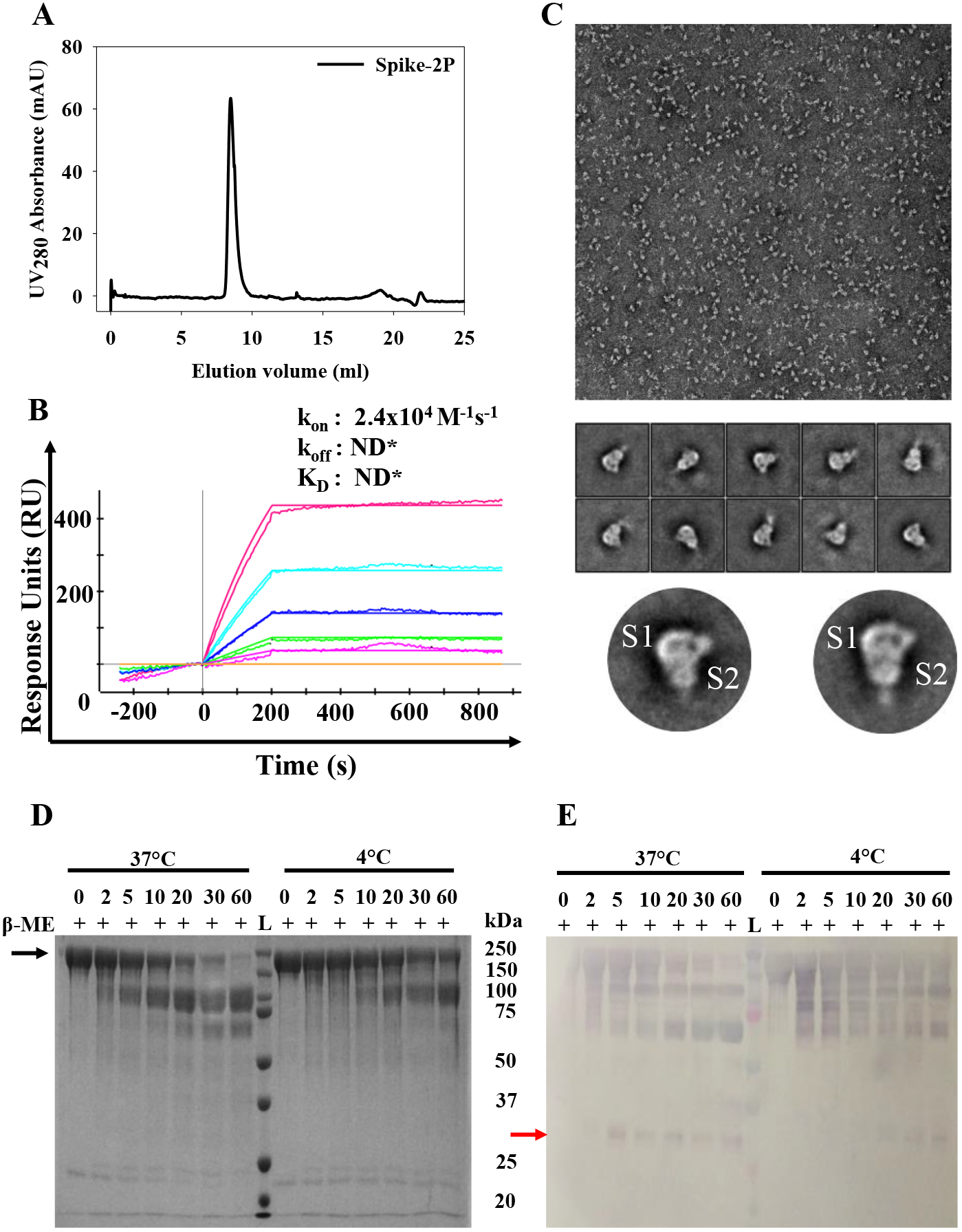
Characterization of mammalian cell expressed Spike-2P. **A)** Size exclusion chromatography profile of Spike-2P ectodomain with a trimeric peak at ~8.9 mL on S200 10/300GL column calibrated with Biorad gel filtration marker (Cat. No. 1511901) run at flowrate of 0.75 mL/min with PBS (pH 7.4) as mobile phase. **B)** SPR binding sensorgrams of *Expi293F* purified Spike-2P with immobilized ACE2-hFc. The concentrations of Spike-2P analyte used are 146 nM, 73 nM, 36.5 nM, 18 nM, 9 nM. **C)** Negative staining EM images of Spike-2P protein. TEM images indicate that the sample is homogeneous and monodisperse. Representative 2D reference free class averages of Spike-2P protein. Well-defined class averages indicate that the Spike-2P sample has a stable and ordered structure and enlarged views of two class averages show the S1 and S2 subunits of spike protein. **D)** SDS-PAGE Coomassie stained gel following limited proteolysis of purified Spike-2P by TPCK treated trypsin (RBD:TPCK Trypsin=50:1) at 4°C and 37°C. **E)** Western blot following limited proteolysis of purified Spike-2P by TPCK treated trypsin (RBD:TPCK Trypsin=50:1) at 4°C and 37°C, probed by α-mRBD guinea pig sera. The red arrow denotes the expected position of the RBD.

Maintaining a proper cold chain during mass vaccination programs can be challenging in low and middle income countries (28, 29). The aggregation state of mRBD and Spike-2P proteins was unchanged upon storage at 4°C, freeze thaw and hour-long 37 °C incubation (Supplementary Figure 2A, 2B). mRBD and Spike were also incubated at various temperatures both in PBS buffer for 60 minutes and for 90 minutes in the lyophilized state. Protein conformational integrity was then assessed in ACE2 binding experiments using SPR. In PBS, mRBD is 20 °C more stable to thermal stress compared to Spike-2P (Figure 4A, 4B). Remarkably, lyophilized mRBD was stable to exposure of temperatures as high as 100 °C whereas Spike-2P rapidly lost activity at temperatures above 50 °C both in solution and in the lyophilized state (Figure 4C, 4D). In solution, mRBD thermal unfolding was highly reversible in contrast to Spike-2P, as assessed by repetitive equilibrium thermal unfolding (Figure 4 E, 4F). mRBD had identical thermal stability profiles before and after lyophilization (Supplementary Figure 2C). mRBD was also resistant to longer time (16 hour) thermal stress and showed only small changes in the thermal unfolding profile when incubated for this time at temperatures up to 70 °C in the lyophilized state, and up to 37 °C in pH 7.0 buffer (Figure 4G, 4H, Supplementary Figures 2D, 2E). Even after storage at 37 °C for four weeks in the lyophilized state, the thermal stability as well the Ace2 binding of the protein were unaffected (Figure 4G, 4H)

**Figure 4:**
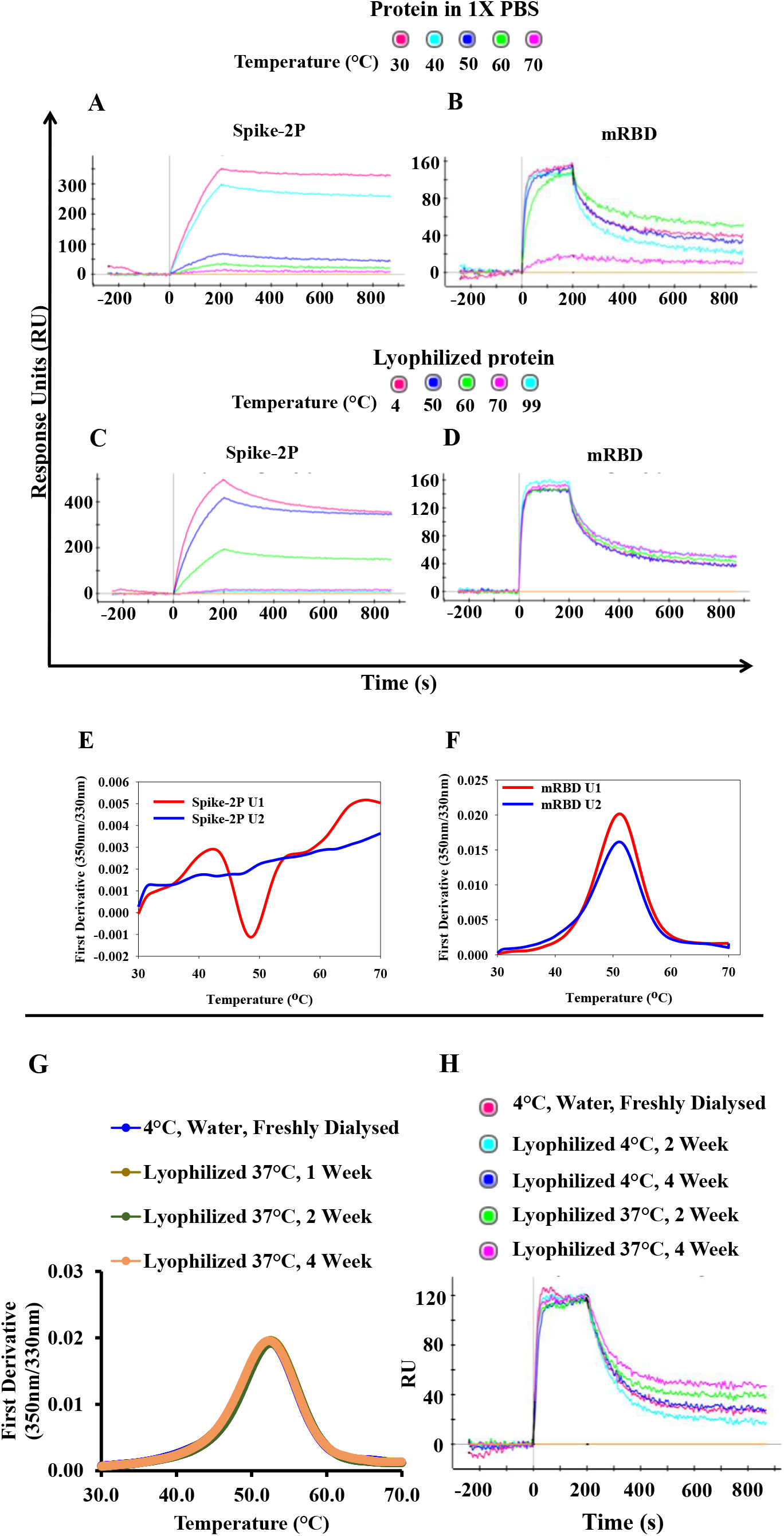
RBD and Spike protein functionality upon subjecting to thermal stress. SPR sensorgrams of ACE2 binding by **A), B)** protein in 1X PBS, subjected to thermal stress for 60 minutes **C), D)** Lyophilized protein subjected to thermal stress for 90 minutes. 100 nM of Spike-2P and mRBD were used as analytes. **E), F)** Equilibrium thermal unfolding measured using nanoDSF for Spike-2P and mRBD. The initial and repeat unfolding scans are in red (U1) and blue (U2) respectively. **G)** Equilibrium thermal unfolding measured using nanoDSF for lyophilized mRBD subjected to 37°C incubation for up to four weeks. **H)** SPR sensorgrams of ACE2 binding by lyophilized mRBD incubated at 4°C, 37°C for up to four weeks. 100nM of mRBD in 1XPBS was used as analyte.

### AddaVax™ adjuvanted RBD elicits neutralizing antibodies in guinea pigs, functionally blocking the receptor binding motif

Guinea pigs are a widely used, outbred animal model for respiratory infectious diseases and display disease susceptibility and immune responses that are more similar to humans than the mouse model (30). Guinea pigs have also been used to evaluate other COVID-19 vaccine candidates (20, 31, 32). Guinea pigs were immunized with mammalian cell expressed mRBD protein adjuvanted with AddaVax™. AddaVax is a squalene-based oil-in-water emulsion that is a mimetic of MF59. MF59 has an extensive safety record and has been used in millions of people in the context of adjuvanted influenza vaccines (33). Animals were primed at day 0 and boosted at day 21 with bleeds at day −1 (Pre-Bleed), day 14 and day 35.

The end point ELISA titers to self-antigen ranged from 1:6400 to 1:102400 after the second immunization in individual animals (Figure 5A). To further confirm and extend these results, the study was repeated with the inclusion of two additional groups immunized with *Pichia* expressed RBD and mammalian cell expressed Spike-2P in addition to the mRBD (Figure 5A, 5B). Results with the mRBD were consistent in both studies (Figure 5A). *Pichia* expressed RBD was as immunogenic as the mRBD in terms of self-titers (Supplementary Figure 3A, 3B) but sera reacted poorly with mRBD and Spike-2P. Several studies have now shown that ACE2 competition titers and neutralizing antibody titers are highly correlated (22, 34). Hence serum competition assays were carried out (Figure 5C). Endpoint neutralization titers with replicative virus were measured using cytopathic effect (CPE) as a readout for infection (Figure 5D) and found to range from 160-1280. Surprisingly, the pRBD sera were non-neutralizing and poorly cross-reactive with mRBD and Spike-2P proteins (Figure 5A, 5C, 5D), presumably because of hyperglycosylation of the *Pichia* expressed protein. In Spike-2P immunized animals titers were more variable than in mRBD immunized animals though the difference in neutralization titers did not approach statistical significance. A potential advantage of using the spike as an immunogen is that it contains neutralization epitopes outside the RBD, including in the NTD (35). We therefore probed the Spike-2P sera for NTD titers using a mammalian cell expressed NTD construct. However, all spike sera had NTD endpoint ELISA titers less than 100. mRBD elicited serum neutralization titers that compared favorably with those observed with several vaccine modalities in a variety of different organisms including guinea pigs (Figure 5E) (18, 19, 21, 36–41)(42). A recent study compared titers elicited by an inactivated virus vaccine formulation (BBIBP-CorV, Figure 5E) in mice, rats, guinea pigs and non-human and non-human primates, the data show close consistency across all the different animal models (31).

**Figure 5:**
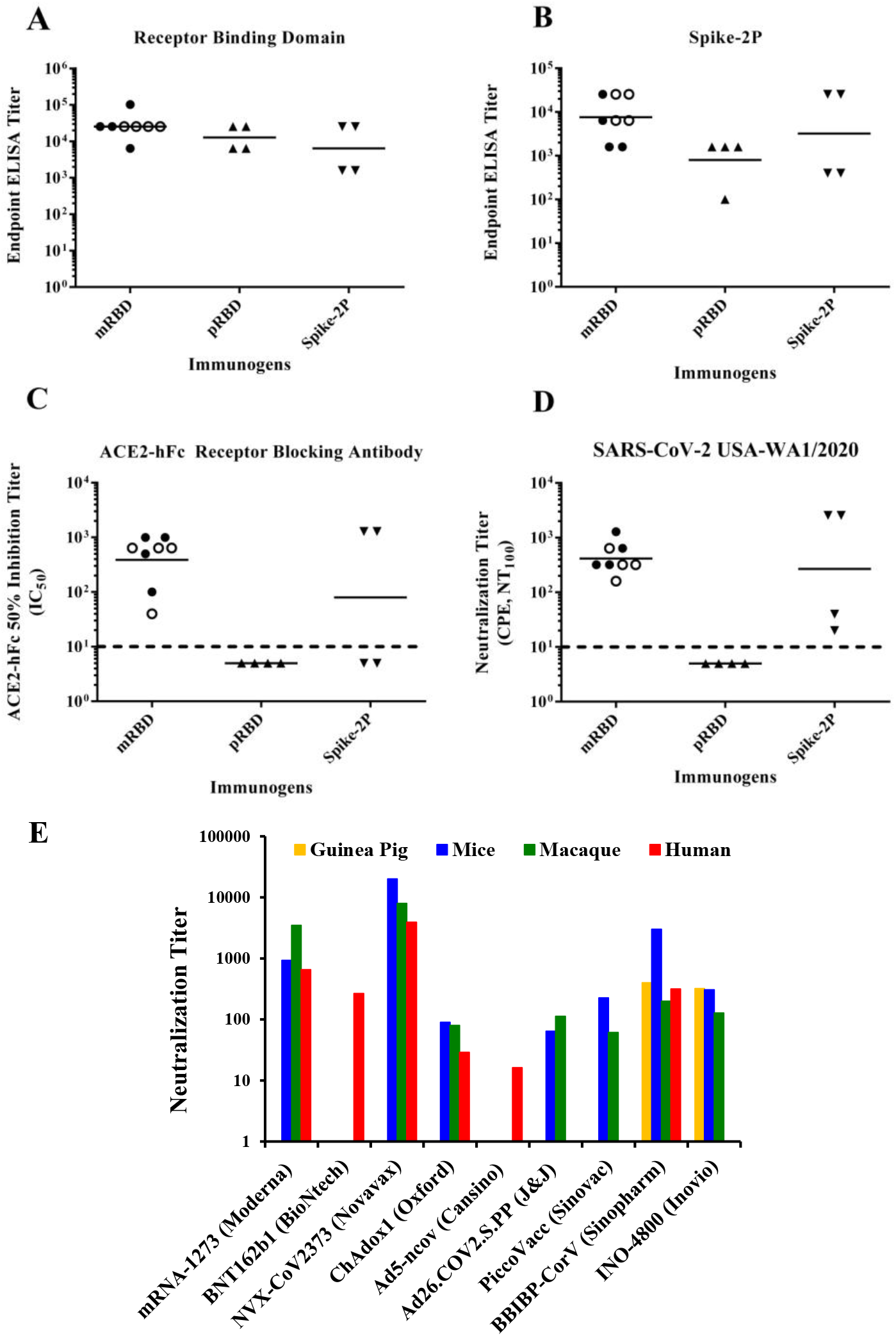
Comparative immunogenicity data. **A-D)** Guinea Pig serum titers obtained after two immunizations with AddaVax™ formulated immunogens. **A), B)** ELISA endpoint titer against mammalian cell expressed RBD and Spike ectodomain respectively. **C)** 50% Inhibitory titers of ACE2 receptor competing antibodies from animals immunized with mRBD, pRBD and Spike-2P respectively. Competition titers below 10 are uniformly assigned a value of 5. The dashed line represents the value 10. **D)** Endpoint neutralization titers in a cytopathic effect (CPE) assay against infectious SARS-CoV-2, Isolate USA-WA1/2020. The dashed line represents the value 10. (•)Sera from the first batch of animals immunized sera with mRBD, (◦) second batch of animals immunized with mRBD, (▴) animals immunized with pRBD, (▾) immunized with mammalian expressed Spike-2P. **E)** Live virus neutralization titers for various vaccine candidates in mice, macaques and humans. mRNA-1273: mRNA vaccine expressing full length Spike-2P protein assayed by PRNT (36, 37, 40). BNT162b1: nucleoside modified mRNA vaccine expressing RBD subunit fused to T4 Fibritin derived Foldon trimerization domain assayed by PRNT(17). NVX-CoV2373: Full length Spike-2P adjuvanted protein vaccine assayed by CPE(22, 43). ChAdOx1 nCoV-19: Replication-deficient Chimpanzee Adenovirus vector expressing spike protein assayed by PRNT and Marburg VN(18, 38, 39). Ad5-ncov: Replication-defective Adenovirus type 5 vector expressing spike protein assayed by CPE(19). Ad26.COV2.S: Replication-defective Adenovirus type 26 vector expressing spike protein assayed by PRNT (41,44). PiccoVacc: Chemically inactivated SARS-CoV-2 virus vaccine assayed by CPE (21). BBIBP: Chemically inactivated SARS-CoV-2 virus vaccine assayed by CPE (31, 42). INO-4800: DNA vaccine expressing full length Spike-2P protein assayed by CPE (20). Macaque data in INO-4800 is obtained by a SARS-CoV-2 pseudovirus neutralization assay (45).

## Discussion

The majority of SARS-CoV-2 vaccine candidates currently in clinical testing use either full length spike or the corresponding ectodomain as antigen, and most involve relatively new nucleic acid or viral vector modalities that have not been tested in large scale immunizations. There is an obvious need for highly expressed, stable, low-cost and efficacious subunit vaccine formulations and for side-by-side comparisons of different candidates. In the present study we characterized the comparative yield, stability and immunogenicity of mammalian cell and *Pichia* expressed RBD, as well as mammalian cell expressed stabilized Spike-2P protein. All three candidates were successfully expressed, properly folded and immunogenic. The data clearly indicate mammalian cell expressed, glycan engineered RBD to be the best of the three immunogens, displaying reversible thermal unfolding and exceptionally high thermal tolerance and stability to storage at 37 °C upto at least four weeks, a very important attribute for deployment in low resource settings. The ability of the mRBD to elicit neutralizing antibodies was comparable to that of the spike ectodomain as also seen in a recent rabbit immunogenicity study with a different RBD derivative comprising residues 319-541(23). The RBD fragment could also be expressed at high yield in the microbial host *Pichia pastoris*, and was properly folded, stable and immunogenic. Interestingly, an alhydrogel adjuvanted formulation of a related SARS-CoV-1 RBD construct was recently shown to be immunogenic and protect mice from SARS-CoV-1 challenge (46). Unfortunately, in the present study when pRBD was used as an immunogen, the elicited antibodies were poorly reactive with either the mammalian cell expressed RBD or the corresponding Spike ectodomain. Further, they failed to block binding of RBD to the ACE2 receptor, suggesting that further alterations to the *Pichia* expressed sequence or adjuvant, use of an alternative *Pichia* strain, or optimization of growth/fermentation conditions are required before it can be used as an effective immunogen. Recently, various RBD derived subunit vaccine candidates have been tested for immunogenicity employing varying fragment lengths, fusion adaptors (Fc, dimers), and adjuvants. No antibody dependent enhancement of infection, immunopathologies, or Th2 bias have been observed with the SARS-CoV-2 RBD subunit derivatives examined so far (46–49). Three independent studies used RBD-Fc fusions with one study using RBD residues 331-527, another used RBD-Fc from Sino Biologicals (residues not mentioned) and a third used a full-length S1-Fc fusion (residues 14-685) reporting viral neutralizing antibody titers of ~100-400, 1280 and NT_50_ derived from pseudoviral neutralizations of 378 respectively (47, 48, 50). One study employed a week long intra-peritoneal immunization regime that is difficult to implement in large-scale human vaccination programs (50). The other studies utilizing RBD-Fc and S1-Fc (47, 48), employed Freunds adjuvant, again not used in human vaccinations. For the present mRBD formulation both the IC_50_ values in the ACE2 competition assay and the viral neutralization titers were about 2% of the corresponding ELISA endpoint titers, suggesting that a significant fraction of the elicited antibodies are neutralizing. Oligomerization and nanoparticle display strategies have proven to induce appreciably higher neutralizing antibody titers than corresponding monomers, this could be potentially be exploited with our mRBD construct in future studies (49, 51, 52). However the effect of these modifications, as well as the exact choice of chain termini which differ between the various RBD constructs, on thermotolerance remain to be studied. Additionally, display on a heterologous scaffold will likely elicit significant antibody titers against the scaffold as well as the displayed immunogen, which might pose regulatory challenges. An mRNA vaccine encoding a longer RBD fragment (319-541) elicited approximately comparable neutralization titers in mice and macaques to those observed in the present study. In the same study, a corresponding luciferase reporter mRNA formulation was shown to tolerate 37 °C incubation for a week with ~13% loss in activity (53). Multiple studies employing a variety of vaccine formulations and modalities have now demonstrated that SARS-CoV-2 viral neutralization titers in small animals, including mice and guinea pigs, are predictive of immunogenicity in macaques and humans (Figure 5E) (17, 18, 40, 46–49, 54, 19, 20, 22, 23, 36–39). Despite promising immunogenicity in several cases, all of the above liquid vaccine formulations were either refrigerated or frozen prior to use. In contrast, the mRBD described above can be stored lyophilized without refrigeration for at least four weeks, and is also tolerant to transient high temperature exposure. In future studies the present formulation will be tested for its ability to confer protection against challenge in an appropriate model, following which it can be advanced to clinical development. It will also be valuable to examine if other RBD protein formulations with different chain termini are similarly thermotolerant.

## Materials and Methods

### SARS-CoV-2 RBD, NTD, Spike ectodomain and antibody expression constructs

Two fragments of the SARS-CoV-2 Spike protein (S) (accession number YP_009724390.1) consisting of the receptor binding domain (RBD) residues 331-532 with an N-terminal glycosylation site and 332-532 with deletion of the N-terminal glycan site deletion were chosen. Residue N532 was engineered to be glycosylated by introducing an NGS motif at the C-termini of the RBD into both immunogen sequences. The resulting sequences with an HRV-3C precision protease cleavage site linked to a 10xHistidine tag by GS linker was codon optimized for human cell expression were expressed under control of the CMV promoter with a tPA signal sequence for efficient secretion. The clones were named m331RBD (331-532) and mRBD (332-532). Identical RBD amino acid sequences to those described above, codon optimized for *Pichia pastoris* expression were cloned into the AOX1 promoter containing vector pPICZalphaA, containing a MATalpha signal sequence for efficient secretion. The resulting clones were named p331RBD (expressing RBD 331-532) and pRBD (expressing RBD 332-532). In the present study, only pRBD was utilized, based on expression data for the corresponding mammalian and insect cell expression clones. A spike N-terminal domain construct (NTD) (residues 27-309 with and L296E mutation) under control of the CMV promoter with a tPA signal sequence was also designed. A spike construct, encoding a stabilized ectodomain with two Proline mutations (Spike-2P) optimized for mammalian cell expression was obtained from the VRC, NIH (5). Genes for the heavy and light chain of the CR3022 antibody were obtained from Genscript (USA) and cloned into the pcDNA3.4 vector

### Purification of recombinant proteins expressed in Expi293F cells

Transfections were performed according to the manufacturer’s guidelines (Gibco, Thermofisher). Briefly, one day prior to transfection, cells were passaged at a density of 2×10^6^cells/mL. On the day of transfection, cells were diluted to 3.0×10^6^cells/mL. Desired plasmids (1μg plasmid per 1mL of Expi293F cells) were complexed with ExpiFectamine293 (2.7μL of ExpiFectamine293 per 1 μg of plasmid) and transiently transfected into Expi293F cells. Post 16hr, Enhancer 1 and Enhancer 2 were added according to the manufacturer’s protocol. Five days post transfection, culture supernatant was collected, proteins were affinity purified by immobilized metal affinity chromatography (IMAC) using Ni Sepharose 6 Fast flow resin (GE Healthcare). Supernatant was two-fold diluted with 1xPBS (pH 7.4) bound to a column equilibrated with PBS (pH7.4). A ten-column volume wash of 1xPBS (pH7.4), supplemented with 25mM imidazole was performed. Finally, the bound protein was eluted with a gradient of 200-500mM imidazole in PBS (pH 7.4). The eluted fractions were pooled, and dialysed thrice using a 3-5kDa (MWCO) dialysis membrane (40mm flat width) (Spectrum Labs) against PBS (pH 7.4). Protein concentration was determined by absorbance (A_280_) using NanoDrop™2000c with the theoretical molar extinction coefficient calculated using the ProtParam tool (ExPASy).

### Purification of recombinant protein expressed in *Pichia pastoris*

20μg of pRBD vector was linearized with the PmeI enzyme by incubating at 37°C overnight (NEB, R0560). Enzyme was inactivated (65°C, 15min) prior to PCR purification of the linearized product (Qiagen, Germany). 10μg of linearized plasmid was transformed into *Pichia pastoris* X-33 strain by electroporation as per the manufacturers protocol (Thermo Fisher). Transformants were selected on Zeocin containing YPDS plates at a Zeocin concentration of 2mg/mL (Thermo Fisher Scientific) after incubation for 3 days at 30°C.

10 colonies from the YPDS plate were picked and screened for expression by inducing with 1% methanol, fresh methanol was added every 24 hrs. Shake flasks (50mL) containing 8mL BMMY media (pH 6.0) each were used for growing the cultures for up to 120 hrs maintained at 30°C, 250rpm. The expression levels were monitored by dot blot analysis with anti-His tag antibodies. The colony showing the highest expression level was then chosen for large scale expression.

Larger scale cultures were performed in shake flasks by maintaining the same volumetric ratio (flask: media) as the small scale cultures. The expression levels were monitored every 24 hrs using sandwich-ELISA.

Cultures were harvested by centrifuging at 4000g and subsequent filtering through a 0.45μm filter (Sartorius). The supernatant was bound to pre-equilibrated Ni Sepharose 6 Fast flow resin (GE Healthcare). The beads were washed with 1xPBS (pH 7.4), supplemented with 150mM NaCl and 20mM imidazole. Finally, the His tagged pRBD protein was eluted in 1xPBS (pH 7.4) supplemented with 150mM NaCl and 300mM imidazole. The eluted fractions were checked for purity on a SDS-PAGE. Following his, appropriate fractions were pooled and dialyzed against 1x PBS (pH7.4) to remove imidazole.

### Purification of recombinant protein from *E. coli*

The *E. coli* expression construct, eInCV01R consisted of residues 331-532 of the RBD expressed under control of the T7 promoter with an N-terminal His tag in the vector pET15b. eInCV01R was transformed in both *E. coli* BL21(DE3) (Novagen) and *E. coli* SHuffle T7 cells (NEB C3029H). Following cell growth in Terrific Broth and induction with 1 mM IPTG at an OD600 of 1 at either 30 or 37□C, cells were grown for ten hours. Expression was seen in the insoluble and soluble fractions in these two strains respectively. Following cell lysis of the SHuffle cells, protein was purified using Ni-NTA chromatography with a yield of about 1 mg/liter. The protein was aggregation prone and failed to bind ACE2-hFc. In the case of BL21(DE3), following cell lysis, inclusion bodies were solubilized in buffer containing 7M Guanidine Hydrochloride and 10mM mercaptoethanol. Protein was purified using Ni-NTA chromatography under denaturing conditions. Protein was diluted into refolding buffer containing 0.4 M L -Arginine, 100 mM Tris-HCl (pH 8.0), 2.0 mM EDTA (pH 8.0), 5.0 mM L-glutathione reduced, 0.5 mM L-glutathione oxidized, but precipitated. Refolding in the absence of redox buffer was also unsuccessful.

### Tag removal

The His tag was removed by subjecting proteins to digestion with HRV-3C protease (Protein: HRV-3C = 50:1) in PBS (pH 7.4) buffer and incubating at 4°C, 16 hrs. The untagged protein (containing C-terminal sequence: LEVLFQ) was separated from the remaining tag protein and protease by immobilized metal affinity chromatography (IMAC) using Ni Sepharose 6 Fast flow resin (GE Healthcare). The unbound tag-free protein was collected and protein concentration was determined by absorbance (A_280_) using NanoDrop™2000c with the theoretical molar extinction coefficient calculated using the ProtParam tool (ExPASy).

### SDS-PAGE and western blot analysis

SDS-PAGE was performed to estimate the purity of the recombinant proteins. Protein samples were denatured by boiling with sample buffer containing SDS, with or without DTT. For western blotting, following SDS-PAGE, proteins were electrophoretically transferred onto an Immobilon-P membrane (Millipore). After transfer, the membrane was blocked with 3% non-fat milk. The membrane was washed with PBST (1xPBS with 0.05% Tween-20) and incubated with antisera raised against mRBD in guinea pig (1:100). Following this blot was washed and incubated with α-guinea pig ALP conjugated antibody (Sigma) at 1:5000. After washing with 1xPBST, blot was developed by BCIP/NBT Liquid substrate system (Sigma).

### Size exclusion chromatography (SEC)

A Superdex-200 10/300GL analytical gel filtration column (GE healthcare) equilibrated in 1xPBS (pH 7.4) buffer was used SEC profiles were obtained using a Biorad NGC chromatography system. The Area under the curve (AUC) was calculated using the peak integrate tool in the Evaluation platform for various peaks from each run

### nanoDSF studies

Equilibrium thermal unfolding experiments of m331RBD (−10xHis tag), mRBD (−10xHis tag), pRBD (−10xHis tag) and Spike-2P were carried out using a nanoDSF (Prometheus NT.48). Two independent measurements were carried out in duplicate with 10-44 μM of protein in the temperature range of 15-95 °C at 40-80% LED power and initial discovery scan counts (350nm) ranging between 5000 and 10000. In all cases, lyophilized protein was redissolved in water, prior to DSF.

### SPR-binding of immobilized ACE2-hFc / CR3022 to Spike-2P and RBD derivatives as analytes

ACE2-hFc and CR3022 neutralizing antibody binding studies with the various RBD derivatives purified from different expression platforms were carried out using the ProteOn XPR36 Protein Interaction Array V.3.1 from Bio-Rad. Activation of the GLM sensor chip was performed by reaction with EDC and sulfo-NHS (Sigma). Protein G (Sigma) at 10 μg/mL was coupled in the presence of 10mM sodium acetate buffer pH 4.5 at 30 μl/min for 300 seconds in various channels. The Response Units for coupling Protein G were monitored till ~3500-4000 RU was immobilized. Finally, the excess sulfo-NHS esters were quenched using 1M ethanolamine. Following this, ~1000 RU of ACE2-hFc or CR3022 was immobilized on various channels at a flow rate of 5 μg/mL for 100 seconds leaving one channel blank that acts as the reference channel. mRBD, pRBD and Spike-2P were passed at a flow rate of 30 μL/min for 200 seconds over the chip surface, followed by a dissociation step of 600 seconds. A lane without any immobilization was used to monitor non-specific binding. After each kinetic assay, the chip was regenerated in 0.1 M Glycine-HCl (pH 2.7) (in the case of the ACE2-hFc assay) and 4M MgCl_2_ (in case of the CR3022 binding assay). The immobilization cycle was repeated prior to each kinetic binding assay in case of ACE2-hFc. Various concentrations of the mRBD (−10xHis tag) (100 nM, 50 nM, 25 nM, 12.5 nM, 6.25 nM), pRBD (−10xHis tag) (100 nM, 50 nM, 25 nM) and Spike-2P (−8xHis tag) (146 nM, 73 nM, 36.5 nM, 18.2 nM, 9.1 nM) in 1x PBST were used for binding studies. The kinetic parameters were obtained by fitting the data to a simple 1:1 Langmuir interaction model using Proteon Manager.

### SPR-binding of immobilized ACE2-hFc to thermal stress subjected mammalian RBD/ Spike-2P as analytes

Mammalian RBD/Spike-2P protein at concentration of 0.2 mg/ml in either 1X PBS or as lyophilized protein was subjected to thermal stress by incubation at the desired temperature in a thermal cycler for sixty or ninety minutes respectively. Following this, lyophilized protein was resuspended in water and SPR binding assay as described above was performed to assess the binding response using 100 nM of the thermally stressed protein.

### Limited Proteolysis

An isothermal limited proteolysis assay was carried out for mRBD, pRBD and Spike-2P using TPCK-Trypsin at 4°C and 37°C. Substrate proteins were dialyzed in autoclaved water (MQ) and reconstituted in the digestion buffer (50 mM Tris, 1 mM CaCl_2_ (pH 7.5)). ~100μg of each protein was subjected to proteolysis with 2 μg of TPCK-trypsin (TPCK Trypsin: Vaccine candidate =1:50) incubated at two different temperatures 4 °C and 37 °C with equal volumes of sample drawn at various time points 0, 2, 5, 10, 20, 30 and 60 minutes respectively. The reaction was quenched by SDS-PAGE loading buffer and incubation at 95 °C and analysed by SDS-PAGE.

### Guinea Pig Immunizations

Groups of four, female, Hartley strain guinea pigs, (6-8 weeks old, approximately weighing 300 g) were immunized with 20 μg of purified antigen protein diluted in 50 μl PBS, (pH 7.4), and mixed with 50 μl of AddaVax™ adjuvant (vac-adx-10) (1:1 v/v Antigen: AddaVax™ ratio per animal/dose) (InvivoGen, USA). Immunizations were given by intramuscular injection on Day 0 (prime) and 21 (boost). Blood was collected and serum isolated on day −2 (pre-bleed), 14 and 35, following the prime and boost immunization, respectively. All animal studies were approved by the Institutional Animal Ethics committee (IAEC) No. RR/IAEC/72-2019, Invivo/GP/084. Although there were lockdown associated constraints on procurement of animals, group sizes of four animals are often used in comparative immunogenicity assessments (55).

### ELISA-serum binding antibody end point titers

96 well plates were coated with immunized vaccine antigen and incubated for two hours at 25 °C (4 μg/mL, in 1xPBS, 50 μL/well) under constant shaking (300 rpm) on a MixMate thermomixer (Eppendorf, USA). ACE2-hFc protein coating was used as a control for antigen immobilization. Following four washes with PBST (200μl/well), wells were blocked with blocking solution (100 μL, 3% skimmed milk in 1xPBST) and incubated for one hour at 25 °C, 300 rpm. Next, antisera (60μL) starting at 1:100 dilution with four-fold serial dilutions were added, and plates incubated for 1 hour at 25 °C, 300 rpm. Three washes with 1xPBST were given (200 μL of 1xPBST/well). Following this, Rabbit ALP enzyme conjugated to anti-Guinea Pig IgG secondary antibody (diluted 1:5000 in blocking buffer) (50 μL/well) was added and incubated for 1 hour at 25 °C, 300 rpm (Sigma-Aldrich). Subsequently, four washes were given (200 μL of 1xPBST/well). pNPP liquid substrate (50 μL/well) (pNPP, Sigma-Aldrich) was added and the plate was incubated for 30 minutes at 37 °C, 300 rpm. Finally, the chromogenic signal was measured at 405 nm. The highest serum dilution which had a signal above the cut off value (0.02 O.D. at 405 nm) was considered as the endpoint titer for ELISA.

### ACE2-hFc competition ELISA

96 well plates were coated with vaccine antigen and incubated overnight at 25 °C (4 μg/mL in 1x PBS, 50 μl/well) under constant shaking (300 rpm) on a MixMate thermomixer (Eppendorf, USA). Ovalbumin (4 μg/mL in 1x PBS, 50 μL/well) coating was used as negative control for mRBD immobilization. Next, four washes with 1xPBST were given (200 μl/well) and wells blocked with blocking solution (100 μL 3% skimmed milk in 1xPBST) for one hour at 25 °C, 300 rpm. Next, anti-sera (60μL) starting at a dilution of 1:10 in blocking solution, were added to sera competition wells and blocking solution alone added to the control wells. Samples were incubated for 1 hour at 25°C, 300 rpm and three washes with 1xPBST were given (200 μL of 1xPBST/well). An additional blocking step was performed for one hour with blocking solution (100μL) incubated at 25°C, 300rpm. Following this, an excess of ACE2-hFc was added (60 μL at 20μg/mL) and samples incubated for one hour at 25°C, 300rpm. Three washes were given (200 μL of PBST/well). Next, rabbit ALP enzyme conjugated to anti-Human IgG secondary antibody (diluted 1:5000 in blocking buffer) (50 μl/well) was added and samples incubated for 1 hour at 25°C, 300 rpm (Sigma-Aldrich). Four washes were given (200 μL of PBST/well). pNPP liquid substrate (50 μL/well) was added and the plate was incubated for 30 minutes at 37 °C, 300 rpm. Finally, the chromogenic signal was measured at 405 nm. The percent competition was calculated using the following equation:

% competition = [Absorbance (Control)-Absorbance (Sera Dilution)] * 100 /

[Absorbance (Control)].

Where, Absorbance (Control) is the Absorbance at 405nm of ACE2-hFc protein binding to RBD in the absence of sera, Absorbance (Sera dilution) is the absorbance from wells where the serum dilution is incubated with ACE2-hFc protein and mRBD.

### Sandwich ELISA for monitoring RBD expression

4ug/mL ACE2 in 1xPBS pH 7.4 was coated onto ELISA strips (Thermo Fisher) for 1 hr and then blocked with 3% BSA solution (1x PBS) for 1 hr at RT. Samples were diluted in the blocking solution and incubated in the wells for 2 hrs at RT. The wells were incubated with anti-His Antibody (1:10000 dilution) conjugated with Horseradish peroxidase (HRP) enzyme for 1 hr at RT following which the reaction was visualized by adding 50μL of the chromogenic substrate, TMB (Thermo Fisher). The reaction was stopped after 20 mins with 50μL of 1M HCl and the absorbance reading at 450nm was obtained from an ELISA Plate reader. Plates were washed with 1xPBS pH 7.4 after each step.

### Negative Staining sample preparation and visualization by Transmission Electron Microscope

For visualization by Transmission Electron Microscope, Spike-2P sample was prepared by conventional negative staining method. Briefly, Carbon coated Cu grids were glow discharged for 30 seconds and 3.5 μL of sample (0.1 mg/mL) was incubated on the grid for 1 minute. The extra sample and buffer solution was blotted out and negative staining was performed using 1% Uranyl Acetate solution for 30 seconds. Freshly prepared grids were air-dried for 30 minutes. The negatively stained sample was visualized at room temperature using a Tecnai T12 electron microscope equipped with a LaB_6_ filament operated at 120 kV using a low electron dose. Images were recorded using a side-mounted Olympus VELITA (2K□2K) CCD camera using defocus ranging from −1.3 to −1.5 and a calibrated pixel size 2.54 Å/pixel at specimen level.

### Reference-free 2D classification using single-particle analysis

The evaluation of micrographs was done with EMAN 2.1 (56). Around 2500 particles projections were picked manually and extracted using e2boxer.py in EMAN2.1 software. 2D particle projections were binned by 2 using e2proc2d.py. Reference free 2D classification of different projections of particle were performed using simple_prime2D of SIMPLE 2.1 software (57)

### CPE based viral Neutralization assay

Guinea pig sera after two immunizations (prime and boost) along with pre-immune (negative control) samples were heat inactivated prior to the virus neutralization assay by incubating at 56 °C for one hour. SARS-CoV-2 (Isolate: USA-WA1/2020) live virus, 100TCID_50_ in a volume of 50 μL was premixed with various dilutions of the serum and incubated at 37°C for one hour in a 5% CO_2_ incubator. Serial dilutions of the incubated premix of virus-serum was added in duplicates into a 96 well plate containing VeroE6 cells (10^4^/well) and cultured for 48/96 hours. After completion of incubation, the plates were assessed for virus induced cytopathic effect (CPE) and the neutralization titre was considered as the highest serum dilution at which no CPE was observed under the microscope.

### Production of Pseudotyped SARS-CoV-2 and pseudovirus neutralisation assay

The full-length synthetic construct of Spike glycoprotein of SARS-CoV-2 (GenBank: MN908947) was synthesized from Genewiz, UK. The complete coding sequences of the spike genes of SARS-CoV-2 and SARS-CoV (GenBank: AY278491) lacking the endoplasmic retention signal sequence were amplified from either the synthetic construct or cDNA and cloned into pCAGGS expression vector (pCAGGS-SARS-2-S and pCAGGS-SARS-S). Pseudotyped coronaviruses were produced as previously described. Briefly, the plasmids pCAGGS-SARS2-S and pCAGGS-SARS-S were transiently expressed on HEK 293T cells using Polyethylenimine (PEI) (Polysciences, USA). 24 h post-transfection, the cells were infected with VSVΔG/GFP virus, incubated for 1 hour, and cells were washed thrice with 1xPBS and replaced with DMEM medium containing 1% FCS and antibiotics. Pseudotyped GFP expressing coronaviruses were harvested from the cell supernatant 24 hpi and concentrated using Amicon columns (Merck). Then the viruses were titrated in Vero E6 cells and stored at −80 °C. A Pseudovirus neutralization assay was performed as described elsewhere with minor modifications (58, 59). Guinea pig sera obtained after the first immunization were tested at a dilution of 1:10-1:80 for the presence of neutralizing antibodies to SARS-CoV-2 using pseudotyped virus. Briefly, Vero E6 cells (10,000 cells/well) were plated in a 96 well plate (Nunc, Thermo Scientific) the day before the neutralization assay. Two-fold serially diluted sera were prepared in 96-well plates, starting at 1:10 dilution. Pseudotyped SARS-CoV2 was diluted in Dulbecco’s Modified Dulbecco’s Medium (DMEM) supplemented with 1% FBS and penicillin-streptomycin. Next, 50μl pseudotyped SARS-CoV-2 was added in each well of plates, and the plates were incubated 37°C for 1h. SARS-CoV pseudotyped virus and SARS-CoV polyclonal antibodies (that cross-react with SARS-CoV-2) were used as a positive control. Subsequently, serum-pseudovirus mixtures were transferred to a plate containing Vero E6 cells for one hour. Then the cells were washed twice with 1xPBS and once with medium, and cells were grown in fresh DMEM medium followed by incubation in a 5% CO2 environment at 37 °C for 24 hours. The neutralization titer was measured by calculating the percentage of GFP positive cells in each well.

## Acknowledgements

We thank Dr. Neil King for kindly providing the ACE2-hFc fusion protein and Drs. Lynda Stuart, Dr. Harry Kleanthous of the Bill and Melinda Gates foundation for helpful discussions. We thank Dr. Barney Graham for kindly providing the Spike-2P construct. This work was funded by a grant from the Bill and Melinda Gates Foundation funding to RV. The following reagent NR-52281, SARS-Related Coronavirus 2, Isolate USA-WA1/2020 was deposited by the Centers for Disease Control and Prevention and obtained through BEI Resources, NIAID, NIH: SARS-Related Coronavirus 2, Isolate USA-WA1/2020. We also acknowledge funding for infrastructural support from the following programs of the Government of India: DST-FIST, UGC Center for Advanced Study, MHRD-FAST, the DBT-IISc Partnership Program, and of a JC Bose Fellowship from DST to RV. S.K.M acknowledges the support of MHRD-IISc doctoral fellowship. M.B. acknowledges the support of CSIR doctoral fellowship. N. Sivaji is acknowledged for his enthusiastic technical support. RS, SM and SB acknowledge the intramural funding received from THSTI under Translational Research Program grant.

## Author Contributions

R.V., S.K.M., conceptualized the work, designed the studies. S.P., R.V., G.N., planned the animal studies. R. S., P.R., A.U., performed ELISA and ACE2-hFc competition experiments. S.K.M., K.K., S.G., S.P., N.G., P.K. performed mRBD, Ace2 and Spike protein expression and characterization. S.G. performed pRBD protein expression and characterization. M.S.K., performed *E.coli* RBD expression and purification. S.A. expressed and characterized the mRBD 333H mutant with assistance from M.S.K. I.P., S. D. provided the EM data and analysis. S.M., S.B., Ram.S. provided CPE neutralizing antibody assay data. J.J., K.T., V.S.R, performed pseudovirus neutralization assays. M.B. contributed to antibody epitope analyses. S.K.M wrote the manuscript with contributions from each author. S.K.M, R.V., G.N. led the studies and edited the paper along with all co-authors.

---

**Supplementary Figure 1:**
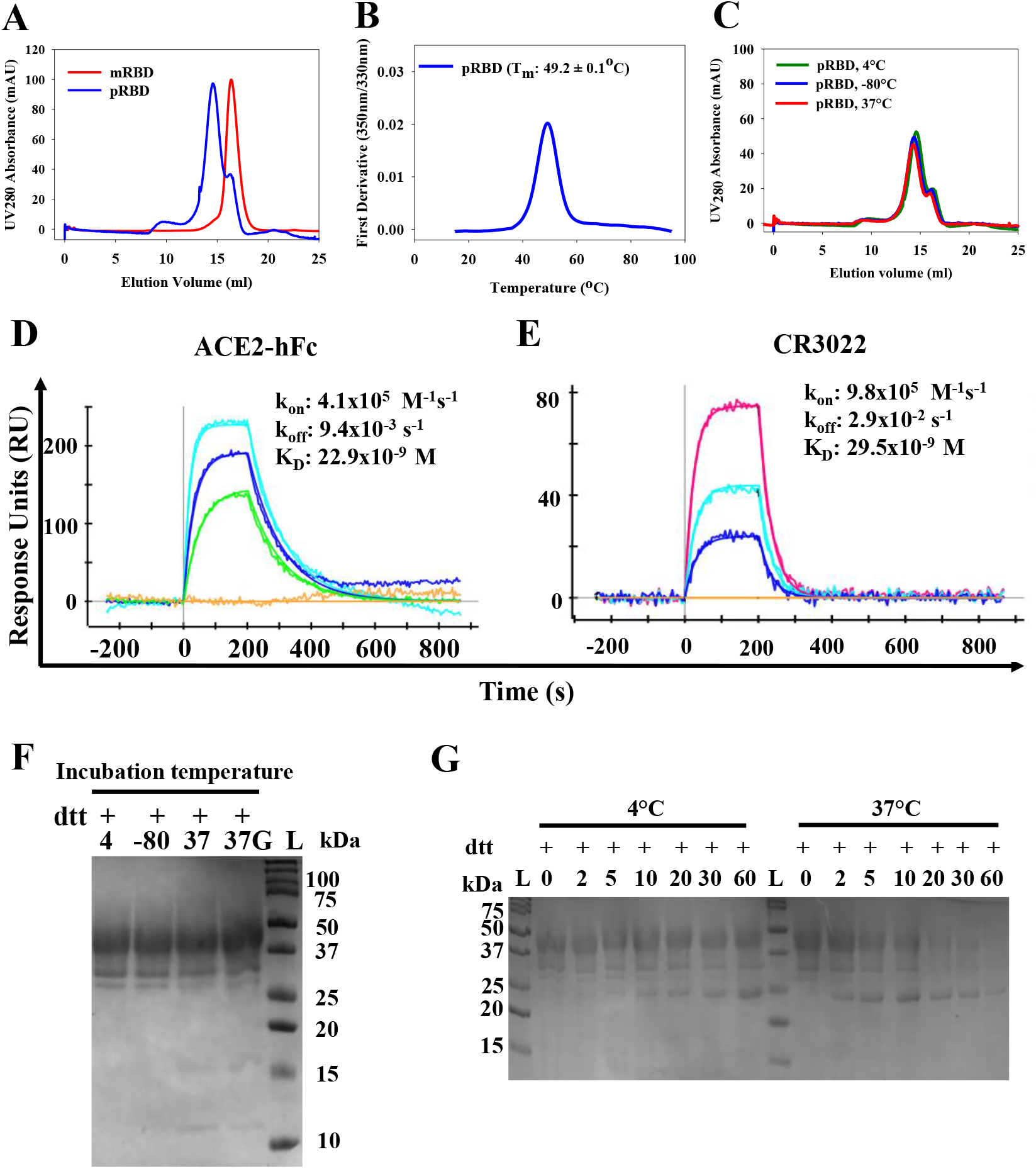
Characterization of pRBD. **A)** Comparison of SEC profiles of *Pichia* and mammalian cell expressed RBD proteins with predominantly monomeric peaks at ~14.5 mL, ~17.0 mL and ~16.3 mL respectively on an S200 10/300GL column calibrated with Biorad gel filtration marker (Cat. No. 1511901) run at flowrate of 0.5mL/min with PBS (pH 7.4) as mobile phase **B)** nanoDSF equilibrium thermal unfolding of pRBD and mRBD. **C)** Size exclusion chromatography profile of pRBD following freeze thaw, incubation at 37°C for one hour and stored overnight at 4 °C. **D)** SPR binding sensorgrams of pRBD to ACE2 receptor. The concentrations of pRBD used as analytes are 100 nM, 50 nM, 25 nM. **E)** SPR binding sensorgrams of pRBD with the neutralizing antibody, CR3022.The concentrations of pRBD used as analytes are 12.5 nM, 6.2 nM, 3.1 nM. **F)** Coomassie stained Reducing SDS-PAGE of pRBD incubated at various temperatures 4-4°C stored protein, −80 −80 °C frozen and thawed protein, 37-protein incubated at 37°C for 1 hour without glycerol, 37G-protein incubated at 37 °C for 1 hour with 5% glycerol. **G)** Limited proteolysis of pRBD with TPCK treated trypsin (RBD:TPCK Trypsin=50:1) at 4°C and 37°C.

**Supplementary Figure 2:**
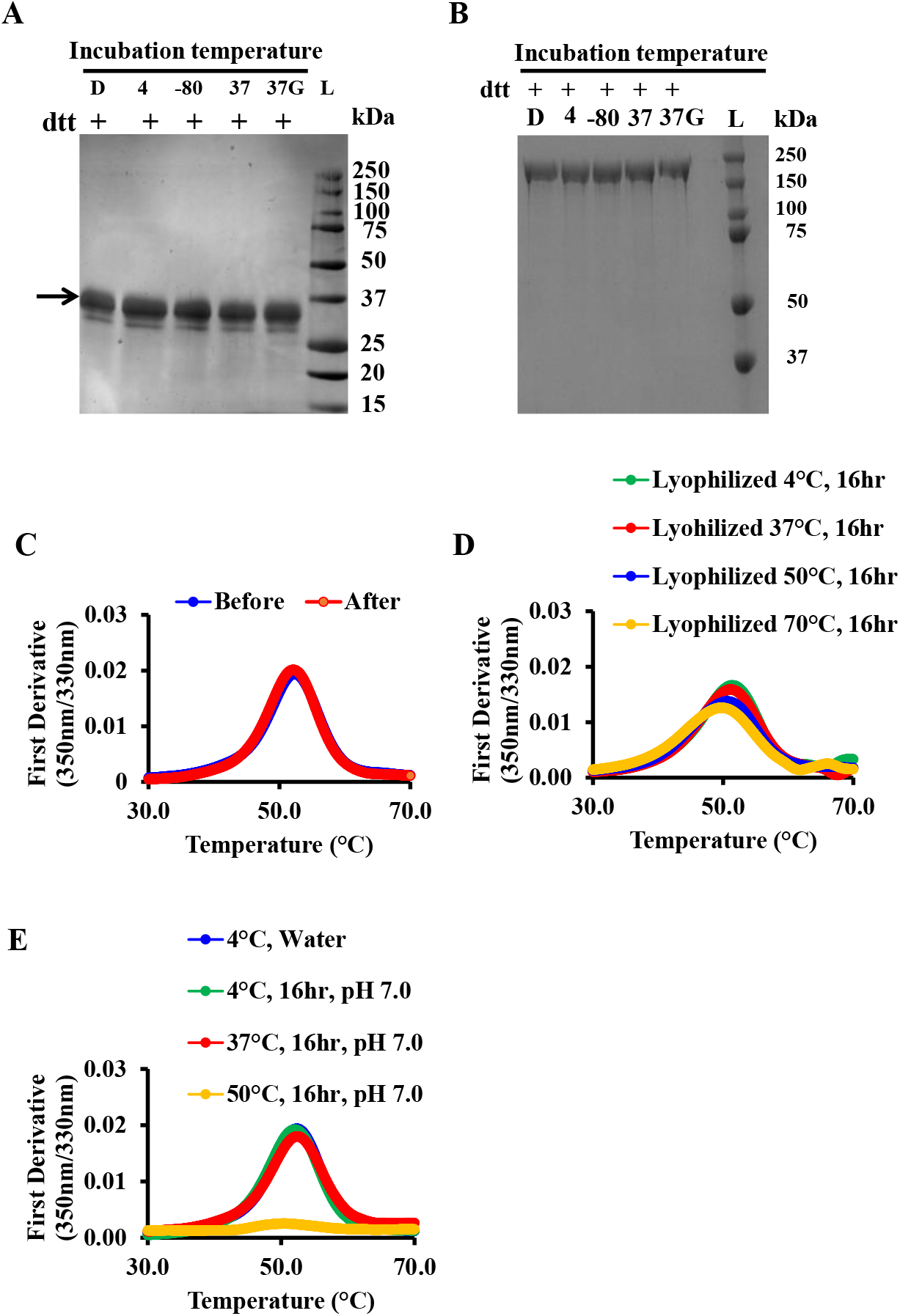
Response of purified mRBD and Spike-2P proteins to various stresses. Coomassie stained reducing SDS-PAGE of **A)** mRBD and **B)** Spike-2P after incubation under different conditions: D-Dialysed and stored overnight at 4 °C, 4-4°C stored protein, −80 °C frozen and thawed protein, 37-protein incubated at 37 °C for 1 hour without glycerol, 37G-protein incubated at 37 °C for 1 hour with 5% glycerol. nanoDSF equilibrium thermal unfolding profiles of **C)** mRBD dialysed against water before (blue) and after (red) lyophilization and resolubilization. **D)** Lyophilized mRBD incubated for 16 hours at 4, 37, 50 and 70 °C and then redissolved in water, prior to nanoDSF. **E)** mRBD in CGH (Citrate,’ Glycine, HEPES (1mM each)) buffer, pH 7, incubated for 16 hours at 4°C, 37 °C and 50 °C. The blue trace in E is identical to that in C and inserted to show that the mRBD thermal unfolding profiles in water and CGH buffer, pH7 are very similar.

**Supplementary Figure 3:**
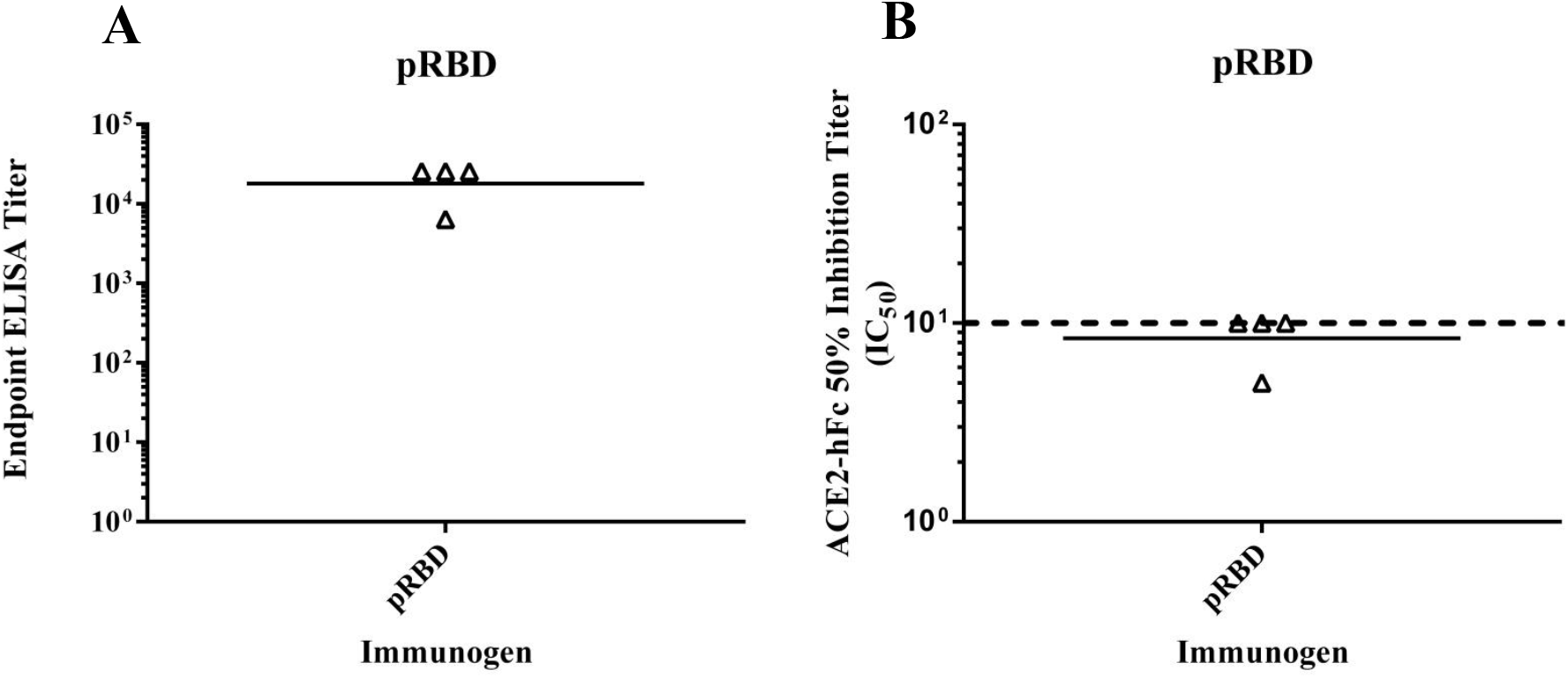
Guinea Pig Serum titers obtained after two immunizations with AddaVax™ formulated pRBD. **A)** ELISA endpoint titer against pRBD **B)** 50% Inhibitory titers of ACE2 receptor competing antibodies from guinea pigs immunized with AddaVax™ adjuvanted pRBD. Competition values below 10 are uniformly assigned a value of five. The dashed line represents the value 10.

## References

1. Chan, J. F.-W., Yuan, S., Kok, K.-H., To, K. K.-W., Chu, H., Yang, J., Xing, F., Liu, J., Yip, C. C.-Y., Poon, R. W.-S., Tsoi, H.-W., Lo, S. K.-F., Chan, K.-H., Poon, V. K.-M., Chan, W.-M., Ip, J. D., Cai, J.-P., Cheng, V. C.-C., Chen, H., Hui, C. K.-M., and Yuen, K.-Y. (2020) A familial cluster of pneumonia associated with the 2019 novel coronavirus indicating person-to-person transmission: a study of a family cluster. Lancet (London, England). 395, 514–523

2. Zhu, N., Zhang, D., Wang, W., Li, X., Yang, B., Song, J., Zhao, X., Huang, B., Shi, W., Lu, R., Niu, P., Zhan, F., Ma, X., Wang, D., Xu, W., Wu, G., Gao, G. F., Tan, W., and China Novel Coronavirus Investigating and Research Team (2020) A Novel Coronavirus from Patients with Pneumonia in China, 2019. N. Engl. J. Med. 382, 727–733

3. WHO (2020) Coronavirus Disease (COVID-19) Dashboard. World Heal. Organ.

4. Bosch, B. J., van der Zee, R., de Haan, C. A. M., and Rottier, P. J. M. (2003) The Coronavirus Spike Protein Is a Class I Virus Fusion Protein: Structural and Functional Characterization of the Fusion Core Complex. J. Virol. 77, 8801–8811

5. Wrapp, D., Wang, N., Corbett, K. S., Goldsmith, J. A., Hsieh, C.-L., Abiona, O., Graham, B. S., and McLellan, J. S. (2020) Cryo-EM structure of the 2019-nCoV spike in the prefusion conformation. Science. 367, 1260–1263

6. Walls, A. C., Park, Y.-J., Tortorici, M. A., Wall, A., McGuire, A. T., and Veesler, D. (2020) Structure, Function, and Antigenicity of the SARS-CoV-2 Spike Glycoprotein. Cell. 181, 281–292.e6

7. Shang, J., Wan, Y., Luo, C., Ye, G., Geng, Q., Auerbach, A., and Li, F. (2020) Cell entry mechanisms of SARS-CoV-2. Proc. Natl. Acad. Sci. 117, 11727–11734

8. Robbiani, D. F., Gaebler, C., Muecksch, F., Lorenzi, J. C. C., Wang, Z., Cho, A., Agudelo, M., Barnes, C. O., Gazumyan, A., Finkin, S., Hägglöf, T., Oliveira, T. Y., Viant, C., Hurley, A., Hoffmann, H.-H., Millard, K. G., Kost, R. G., Cipolla, M., Gordon, K., Bianchini, F., Chen, S. T., Ramos, V., Patel, R., Dizon, J., Shimeliovich, I., Mendoza, P., Hartweger, H., Nogueira, L., Pack, M., Horowitz, J., Schmidt, F., Weisblum, Y., Michailidis, E., Ashbrook, A. W., Waltari, E., Pak, J. E., Huey-Tubman, K. E., Koranda, N., Hoffman, P. R., West, A. P., Rice, C. M., Hatziioannou, T., Bjorkman, P. J., Bieniasz, P. D., Caskey, M., and Nussenzweig, M. C. (2020) Convergent antibody responses to SARS-CoV-2 in convalescent individuals. Nature. 584, 437–442

9. Wang, Q., Zhang, Y., Wu, L., Niu, S., Song, C., Zhang, Z., Lu, G., Qiao, C., Hu, Y., Yuen, K.-Y., Wang, Q., Zhou, H., Yan, J., and Qi, J. (2020) Structural and Functional Basis of SARS-CoV-2 Entry by Using Human ACE2. Cell. 181, 894–904.e9

10. Huo, J., Zhao, Y., Ren, J., Zhou, D., Duyvesteyn, H. M. E., Ginn, H. M., Carrique, L., Malinauskas, T., Ruza, R. R., Shah, P. N. M., Tan, T. K., Rijal, P., Coombes, N., Bewley, K. R., Tree, J. A., Radecke, J., Paterson, N. G., Supasa, P., Mongkolsapaya, J., Screaton, G. R., Carroll, M., Townsend, A., Fry, E. E., Owens, R. J., and Stuart, D. I. (2020) Neutralization of SARS-CoV-2 by Destruction of the Prefusion Spike. Cell Host Microbe. 28, 445–454.e6

11. Pinto, D., Park, Y.-J., Beltramello, M., Walls, A. C., Tortorici, M. A., Bianchi, S., Jaconi, S., Culap, K., Zatta, F., De Marco, A., Peter, A., Guarino, B., Spreafico, R., Cameroni, E., Case, J. B., Chen, R. E., Havenar-Daughton, C., Snell, G., Telenti, A., Virgin, H. W., Lanzavecchia, A., Diamond, M. S., Fink, K., Veesler, D., and Corti, D. (2020) Cross-neutralization of SARS-CoV-2 by a human monoclonal SARS-CoV antibody. Nature. 583, 290–295

12. Rogers, T. F., Zhao, F., Huang, D., Beutler, N., Burns, A., He, W.-T., Limbo, O., Smith, C., Song, G., Woehl, J., Yang, L., Abbott, R. K., Callaghan, S., Garcia, E., Hurtado, J., Parren, M., Peng, L., Ramirez, S., Ricketts, J., Ricciardi, M. J., Rawlings, S. A., Wu, N. C., Yuan, M., Smith, D. M., Nemazee, D., Teijaro, J. R., Voss, J. E., Wilson, I. A., Andrabi, R., Briney, B., Landais, E., Sok, D., Jardine, J. G., and Burton, D. R. (2020) Isolation of potent SARS-CoV-2 neutralizing antibodies and protection from disease in a small animal model. Science (80-.). 10.1126/science.abc7520

13. Shi, R., Shan, C., Duan, X., Chen, Z., Liu, P., Song, J., Song, T., Bi, X., Han, C., Wu, L., Gao, G., Hu, X., Zhang, Y., Tong, Z., Huang, W., Liu, W. J., Wu, G., Zhang, B., Wang, L., Qi, J., Feng, H., Wang, F.-S., Wang, Q., Gao, G. F., Yuan, Z., and Yan, J. (2020) A human neutralizing antibody targets the receptor-binding site of SARS-CoV-2. Nature. 584, 120–124

14. Wu, Y., Wang, F., Shen, C., Peng, W., Li, D., Zhao, C., Li, Z., Li, S., Bi, Y., Yang, Y., Gong, Y., Xiao, H., Fan, Z., Tan, S., Wu, G., Tan, W., Lu, X., Fan, C., Wang, Q., Liu, Y., Zhang, C., Qi, J., Gao, G. F., Gao, F., and Liu, L. (2020) A noncompeting pair of human neutralizing antibodies block COVID-19 virus binding to its receptor ACE2. Science (80-.). 368, 1274–1278

15. Chi, X., Yan, R., Zhang, J., Zhang, G., Zhang, Y., Hao, M., Zhang, Z., Fan, P., Dong, Y., Yang, Y., Chen, Z., Guo, Y., Zhang, J., Li, Y., Song, X., Chen, Y., Xia, L., Fu, L., Hou, L., Xu, J., Yu, C., Li, J., Zhou, Q., and Chen, W. (2020) A neutralizing human antibody binds to the N-terminal domain of the Spike protein of SARS-CoV-2. Science (80-.). 369, 650–655

16. WHO (2020) Draft landscape of COVID-19 candidate vaccines. Who

17. Mulligan, M. J., Lyke, K. E., Kitchin, N., Absalon, J., Gurtman, A., Lockhart, S., Neuzil, K., Raabe, V., Bailey, R., Swanson, K. A., Li, P., Koury, K., Kalina, W., Cooper, D., Fontes-Garfias, C., Shi, P.-Y., Türeci, Ö., Tompkins, K. R., Walsh, E. E., Frenck, R., Falsey, A. R., Dormitzer, P. R., Gruber, W. C., Şahin, U., and Jansen, K. U. (2020) Phase 1/2 study of COVID-19 RNA vaccine BNT162b1 in adults. Nature. 10.1038/s41586-020-2639-4

18. van Doremalen, N., Lambe, T., Spencer, A., Belij-Rammerstorfer, S., Purushotham, J. N., Port, J. R., Avanzato, V., Bushmaker, T., Flaxman, A., Ulaszewska, M., Feldmann, F., Allen, E. R., Sharpe, H., Schulz, J., Holbrook, M., Okumura, A., Meade-White, K., Pérez-Pérez, L., Bissett, C., Gilbride, C., Williamson, B. N., Rosenke, R., Long, D., Ishwarbhai, A., Kailath, R., Rose, L., Morris, S., Powers, C., Lovaglio, J., Hanley, P. W., Scott, D., Saturday, G., de Wit, E., Gilbert, S. C., and Munster, V. J. (2020) ChAdOx1 nCoV-19 vaccination prevents SARS-CoV-2 pneumonia in rhesus macaques. bioRxiv. 10.1101/2020.05.13.093195

19. Zhu, F.-C., Guan, X.-H., Li, Y.-H., Huang, J.-Y., Jiang, T., Hou, L.-H., Li, J.-X., Yang, B.-F., Wang, L., Wang, W.-J., Wu, S.-P., Wang, Z., Wu, X.-H., Xu, J.-J., Zhang, Z., Jia, S.-Y., Wang, B.-S., Hu, Y., Liu, J.-J., Zhang, J., Qian, X.-A., Li, Q., Pan, H.-X., Jiang, H.-D., Deng, P., Gou, J.-B., Wang, X.-W., Wang, X.-H., and Chen, W. (2020) Immunogenicity and safety of a recombinant adenovirus type-5-vectored COVID-19 vaccine in healthy adults aged 18 years or older: a randomised, double-blind, placebo-controlled, phase 2 trial. Lancet. 396, 479–488

20. Smith, T. R. F., Patel, A., Ramos, S., Elwood, D., Zhu, X., Yan, J., Gary, E. N., Walker, S. N., Schultheis, K., Purwar, M., Xu, Z., Walters, J., Bhojnagarwala, P., Yang, M., Chokkalingam, N., Pezzoli, P., Parzych, E., Reuschel, E. L., Doan, A., Tursi, N., Vasquez, M., Choi, J., Tello-Ruiz, E., Maricic, I., Bah, M. A., Wu, Y., Amante, D., Park, D. H., Dia, Y., Ali, A. R., Zaidi, F. I., Generotti, A., Kim, K. Y., Herring, T. A., Reeder, S., Andrade, V. M., Buttigieg, K., Zhao, G., Wu, J.-M., Li, D., Bao, L., Liu, J., Deng, W., Qin, C., Brown, A. S., Khoshnejad, M., Wang, N., Chu, J., Wrapp, D., McLellan, J. S., Muthumani, K., Wang, B., Carroll, M. W., Kim, J. J., Boyer, J., Kulp, D. W., Humeau, L. M. P. F., Weiner, D. B., and Broderick, K. E. (2020) Immunogenicity of a DNA vaccine candidate for COVID-19. Nat. Commun. 11, 2601

21. Gao, Q., Bao, L., Mao, H., Wang, L., Xu, K., Yang, M., Li, Y., Zhu, L., Wang, N., Lv, Z., Gao, H., Ge, X., Kan, B., Hu, Y., Liu, J., Cai, F., Jiang, D., Yin, Y., Qin, C., Li, J., Gong, X., Lou, X., Shi, W., Wu, D., Zhang, H., Zhu, L., Deng, W., Li, Y., Lu, J., Li, C., Wang, X., Yin, W., Zhang, Y., and Qin, C. (2020) Development of an inactivated vaccine candidate for SARS-CoV-2. Science (80-.). 369, 77–81

22. Tian, J.-H., Patel, N., Haupt, R., Zhou, H., Weston, S., Hammond, H., Lague, J., Portnoff, A. D., Norton, J., Guebre-Xabier, M., Zhou, B., Jacobson, K., Maciejewski, S., Khatoon, R., Wisniewska, M., Mottitt, W., Kluepfel-Stahl, S., Ekechukwu, B., Papin, J., Boddapati, S., Wong, C. J., Piedra, P. A., Frieman, M. B., Massare, M. J., Fries, L., Bengtsson, K. L., Stertman, L., Ellingsworth, L. R., Glenn, G., and Smith, G. (2020) SARS-CoV-2 spike glycoprotein vaccine candidate NVX-CoV2373 elicits immunogenicity in baboons and protection in mice. bioRxiv. 10.1101/2020.06.29.178509

23. Ravichandran, S., Coyle, E. M., Klenow, L., Tang, J., Grubbs, G., Liu, S., Wang, T., Golding, H., and Khurana, S. (2020) Antibody signature induced by SARS-CoV-2 spike protein immunogens in rabbits. Sci. Transl. Med. 12, eabc3539

24. Yuan, M., Wu, N. C., Zhu, X., Lee, C.-C. D., So, R. T. Y., Lv, H., Mok, C. K. P., and Wilson, I. A. (2020) A highly conserved cryptic epitope in the receptor binding domains of SARS-CoV-2 and SARS-CoV. Science (80-.). 368, 630–633

25. Zhou, P., Yang, X.-L., Wang, X.-G., Hu, B., Zhang, L., Zhang, W., Si, H.-R., Zhu, Y., Li, B., Huang, C.-L., Chen, H.-D., Chen, J., Luo, Y., Guo, H., Jiang, R.-D., Liu, M.-Q., Chen, Y., Shen, X.-R., Wang, X., Zheng, X.-S., Zhao, K., Chen, Q.-J., Deng, F., Liu, L.-L., Yan, B., Zhan, F.-X., Wang, Y.-Y., Xiao, G.-F., and Shi, Z.-L. (2020) A pneumonia outbreak associated with a new coronavirus of probable bat origin. Nature. 579, 270–273

26. Lan, J., Ge, J., Yu, J., Shan, S., Zhou, H., Fan, S., Zhang, Q., Shi, X., Wang, Q., Zhang, L., and Wang, X. (2020) Structure of the SARS-CoV-2 spike receptor-binding domain bound to the ACE2 receptor. Nature. 581, 215–220

27. Chen, W.-H., Du, L., Chag, S. M., Ma, C., Tricoche, N., Tao, X., Seid, C. A., Hudspeth, E. M., Lustigman, S., Tseng, C.-T. K., Bottazzi, M. E., Hotez, P. J., Zhan, B., and Jiang, S. (2014) Yeast-expressed recombinant protein of the receptor-binding domain in SARS-CoV spike protein with deglycosylated forms as a SARS vaccine candidate. Hum. Vaccin. Immunother. 10, 648–658

28. Lee, B. Y., Cakouros, B. E., Assi, T.-M., Connor, D. L., Welling, J., Kone, S., Djibo, A., Wateska, A. R., Pierre, L., and Brown, S. T. (2012) The impact of making vaccines thermostable in Niger’s vaccine supply chain. Vaccine. 30, 5637–43

29. Kristensen, D., Chen, D., and Cummings, R. (2011) Vaccine stabilization: research, commercialization, and potential impact. Vaccine. 29, 7122–4

30. Klein, T. W., Friedman, H., and Widen, R. (1984) Relative potency of virulent versus avirulent Legionella pneumophila for induction of cell-mediated immunity. Infect. Immun. 44, 753–755

31. Wang, H., Zhang, Y., Huang, B., Deng, W., Quan, Y., Wang, W., Xu, W., Zhao, Y., Li, N., Zhang, J., Liang, H., Bao, L., Xu, Y., Ding, L., Zhou, W., Gao, H., Liu, J., Niu, P., Zhao, L., Zhen, W., Fu, H., Yu, S., Zhang, Z., Xu, G., Li, C., Lou, Z., Xu, M., Qin, C., Wu, G., Gao, G. F., Tan, W., and Yang, X. (2020) Development of an Inactivated Vaccine Candidate, BBIBP-CorV, with Potent Protection against SARS-CoV-2. Cell. 182, 713–721.e9

32. WHO (2020) Global animal laboratories capacities to support vaccine and therapeutic evaluation. Who

33. O’Hagan, D. T., Ott, G. S., Nest, G. Van, Rappuoli, R., and Giudice, G. Del (2013) The history of MF59 ® adjuvant: a phoenix that arose from the ashes. Expert Rev. Vaccines. 12, 13–30

34. Abe, K. T., Li, Z., Samson, R., Samavarchi-Tehrani, P., Valcourt, E. J., Wood, H., Budylowski, P., Dupuis, A., Girardin, R. C., Rathod, B., Colwill, K., McGeer, A. J., Mubareka, S., Gommerman, J. L., Durocher, Y., Ostrowski, M., McDonough, K. A., Drebot, M. A., Drews, S. J., Rini, J. M., and Gingras, A.-C. (2020) A simple protein-based SARS-CoV-2 surrogate neutralization assay. bioRxiv. 10.1101/2020.07.10.197913

35. Liu, L., Wang, P., Nair, M. S., Yu, J., Rapp, M., Wang, Q., Luo, Y., Chan, J. F. W., Sahi, V., Figueroa, A., Guo, X. V., Cerutti, G., Bimela, J., Gorman, J., Zhou, T., Chen, Z., Yuen, K.-Y., Kwong, P. D., Sodroski, J. G., Yin, M. T., Sheng, Z., Huang, Y., Shapiro, L., and Ho, D. D. (2020) Potent neutralizing antibodies against multiple epitopes on SARS-CoV-2 spike. Nature. 584, 450–456

36. Corbett, K. S., Edwards, D. K., Leist, S. R., Abiona, O. M., Boyoglu-Barnum, S., Gillespie, R. A., Himansu, S., Schäfer, A., Ziwawo, C. T., DiPiazza, A. T., Dinnon, K. H., Elbashir, S. M., Shaw, C. A., Woods, A., Fritch, E. J., Martinez, D. R., Bock, K. W., Minai, M., Nagata, B. M., Hutchinson, G. B., Wu, K., Henry, C., Bahi, K., Garcia-Dominguez, D., Ma, L., Renzi, I., Kong, W.-P., Schmidt, S. D., Wang, L., Zhang, Y., Phung, E., Chang, L. A., Loomis, R. J., Altaras, N. E., Narayanan, E., Metkar, M., Presnyak, V., Liu, C., Louder, M. K., Shi, W., Leung, K., Yang, E. S., West, A., Gully, K. L., Stevens, L. J., Wang, N., Wrapp, D., Doria-Rose, N. A., Stewart-Jones, G., Bennett, H., Alvarado, G. S., Nason, M. C., Ruckwardt, T. J., McLellan, J. S., Denison, M. R., Chappell, J. D., Moore, I. N., Morabito, K. M., Mascola, J. R., Baric, R. S., Carfi, A., and Graham, B. S. (2020) SARS-CoV-2 mRNA vaccine design enabled by prototype pathogen preparedness. Nature. 10.1038/s41586-020-2622-0

37. Corbett, K. S., Flynn, B., Foulds, K. E., Francica, J. R., Boyoglu-Barnum, S., Werner, A. P., Flach, B., O’Connell, S., Bock, K. W., Minai, M., Nagata, B. M., Andersen, H., Martinez, D. R., Noe, A. T., Douek, N., Donaldson, M. M., Nji, N. N., Alvarado, G. S., Edwards, D. K., Flebbe, D. R., Lamb, E., Doria-Rose, N. A., Lin, B. C., Louder, M. K., O’Dell, S., Schmidt, S. D., Phung, E., Chang, L. A., Yap, C., Todd, J.-P. M., Pessaint, L., Van Ry, A., Browne, S., Greenhouse, J., Putman-Taylor, T., Strasbaugh, A., Campbell, T.-A., Cook, A., Dodson, A., Steingrebe, K., Shi, W., Zhang, Y., Abiona, O. M., Wang, L., Pegu, A., Yang, E. S., Leung, K., Zhou, T., Teng, I.-T., Widge, A., Gordon, I., Novik, L., Gillespie, R. A., Loomis, R. J., Moliva, J. I., Stewart-Jones, G., Himansu, S., Kong, W.-P., Nason, M. C., Morabito, K. M., Ruckwardt, T. J., Ledgerwood, J. E., Gaudinski, M. R., Kwong, P. D., Mascola, J. R., Carfi, A., Lewis, M. G., Baric, R. S., McDermott, A., Moore, I. N., Sullivan, N. J., Roederer, M., Seder, R. A., and Graham, B. S. (2020) Evaluation of the mRNA-1273 Vaccine against SARS-CoV-2 in Nonhuman Primates. N. Engl. J. Med. 10.1056/NEJMoa2024671

38. van Doremalen, N., Lambe, T., Spencer, A., Belij-Rammerstorfer, S., Purushotham, J. N., Port, J. R., Avanzato, V., Bushmaker, T., Flaxman, A., Ulaszewska, M., Feldmann, F., Allen, E. R., Sharpe, H., Schulz, J., Holbrook, M., Okumura, A., Meade-White, K., Pérez-Pérez, L., Bissett, C., Gilbride, C., Williamson, B. N., Rosenke, R., Long, D., Ishwarbhai, A., Kailath, R., Rose, L., Morris, S., Powers, C., Lovaglio, J., Hanley, P. W., Scott, D., Saturday, G., de Wit, E., Gilbert, S. C., and Munster, V. J. (2020) ChAdOx1 nCoV-19 vaccination prevents SARS-CoV-2 pneumonia in rhesus macaques. bioRxiv. 10.1101/2020.05.13.093195

39. Graham, S. P., McLean, R. K., Spencer, A. J., Belij-Rammerstorfer, S., Wright, D., Ulaszewska, M., Edwards, J. C., Hayes, J. W. P., Martini, V., Thakur, N., Conceicao, C., Dietrich, I., Shelton, H., Waters, R., Ludi, A., Wilsden, G., Browning, C., Bialy, D., Bhat, S., Stevenson-Leggett, P., Hollinghurst, P., Gilbride, C., Pulido, D., Moffat, K., Sharpe, H., Allen, E., Mioulet, V., Chiu, C., Newman, J., Asfor, A. S., Burman, A., Crossley, S., Huo, J., Owens, R. J., Carroll, M., Hammond, J. A., Tchilian, E., Bailey, D., Charleston, B., Gilbert, S. C., Tuthill, T. J., and Lambe, T. (2020) Evaluation of the immunogenicity of prime-boost vaccination with the replication-deficient viral vectored COVID-19 vaccine candidate ChAdOx1 nCoV-19. npj Vaccines. 5, 69

40. Jackson, L. A., Anderson, E. J., Rouphael, N. G., Roberts, P. C., Makhene, M., Coler, R. N., McCullough, M. P., Chappell, J. D., Denison, M. R., Stevens, L. J., Pruijssers, A. J., McDermott, A., Flach, B., Doria-Rose, N. A., Corbett, K. S., Morabito, K. M., O’Dell, S., Schmidt, S. D., Swanson, P. A., Padilla, M., Mascola, J. R., Neuzil, K. M., Bennett, H., Sun, W., Peters, E., Makowski, M., Albert, J., Cross, K., Buchanan, W., Pikaart-Tautges, R., Ledgerwood, J. E., Graham, B. S., and Beigel, J. H. (2020) An mRNA Vaccine against SARS-CoV-2 — Preliminary Report. N. Engl. J. Med. 0, NEJMoa2022483

41. Mercado, N. B., Zahn, R., Wegmann, F., Loos, C., Chandrashekar, A., Yu, J., Liu, J., Peter, L., McMahan, K., Tostanoski, L. H., He, X., Martinez, D. R., Rutten, L., Bos, R., van Manen, D., Vellinga, J., Custers, J., Langedijk, J. P., Kwaks, T., Bakkers, M. J. G., Zuijdgeest, D., Rosendahl Huber, S. K., Atyeo, C., Fischinger, S., Burke, J. S., Feldman, J., Hauser, B. M., Caradonna, T. M., Bondzie, E. A., Dagotto, G., Gebre, M. S., Hoffman, E., Jacob-Dolan, C., Kirilova, M., Li, Z., Lin, Z., Mahrokhian, S. H., Maxfield, L. F., Nampanya, F., Nityanandam, R., Nkolola, J. P., Patel, S., Ventura, J. D., Verrington, K., Wan, H., Pessaint, L., Van Ry, A., Blade, K., Strasbaugh, A., Cabus, M., Brown, R., Cook, A., Zouantchangadou, S., Teow, E., Andersen, H., Lewis, M. G., Cai, Y., Chen, B., Schmidt, A. G., Reeves, R. K., Baric, R. S., Lauffenburger, D. A., Alter, G., Stoffels, P., Mammen, M., Van Hoof, J., Schuitemaker, H., and Barouch, D. H. (2020) Single-shot Ad26 vaccine protects against SARS-CoV-2 in rhesus macaques. Nature. 10.1038/s41586-020-2607-z

42. Xia, S., Duan, K., Zhang, Y., Zhao, D., Zhang, H., Xie, Z., Li, X., Peng, C., Zhang, Y., Zhang, W., Yang, Y., Chen, W., Gao, X., You, W., Wang, X., Wang, Z., Shi, Z., Wang, Y., Yang, X., Zhang, L., Huang, L., Wang, Q., Lu, J., Yang, Y., Guo, J., Zhou, W., Wan, X., Wu, C., Wang, W., Huang, S., Du, J., Meng, Z., Pan, A., Yuan, Z., Shen, S., Guo, W., and Yang, X. (2020) Effect of an Inactivated Vaccine Against SARS-CoV-2 on Safety and Immunogenicity Outcomes. JAMA. 324, 951

43. Novavax Inc. Novavax Announces Positive Phase 1 Data for its COVID-19 Vaccine Candidate. Novavax Inc. - IR Site

44. Bos, R., Rutten, L., van der Lubbe, J. E. M., Bakkers, M. J. G., Hardenberg, G., Wegmann, F., Zuijdgeest, D., de Wilde, A. H., Koornneef, A., Verwilligen, A., van Manen, D., Kwaks, T., Vogels, R., Dalebout, T. J., Myeni, S. K., Kikkert, M., Snijder, E. J., Li, Z., Barouch, D. H., Vellinga, J., Langedijk, J. P. M., Zahn, R. C., Custers, J., and Schuitemaker, H. (2020) Ad26-vector based COVID-19 vaccine encoding a prefusion stabilized SARS-CoV-2 Spike immunogen induces potent humoral and cellular immune responses. bioRxiv. 10.1101/2020.07.30.227470

45. Patel, A., Walters, J., Reuschel, E. L., Schultheis, K., Parzych, E., Gary, E. N., Maricic, I., Purwar, M., Eblimit, Z., Walker, S. N., Guimet, D., Bhojnagarwala, P., Doan, A., Xu, Z., Elwood, D., Reeder, S. M., Pessaint, L., Kim, K. Y., Cook, A., Chokkalingam, N., Finneyfrock, B., Tello-Ruiz, E., Dodson, A., Choi, J., Generotti, A., Harrison, J., Tursi, N. J., Andrade, V. M., Dia, Y., Zaidi, F. I., Andersen, H., Lewis, M. G., Muthumani, K., Kim, J. J., Kulp, D. W., Humeau, L. M., Ramos, S., Smith, T. R. F., Weiner, D. B., and Broderick, K. E. (2020) Intradermal-delivered DNA vaccine provides anamnestic protection in a rhesus macaque SARS-CoV-2 challenge model. bioRxiv. 10.1101/2020.07.28.225649

46. Chen, W.-H., Tao, X., Peng, B.-H., Pollet, J., Strych, U., Bottazzi, M. E., Hotez, P. J., Lustigman, S., Du, L., Jiang, S., and Tseng, C.-T. K. (2020) Yeast-Expressed SARS-CoV Recombinant Receptor-Binding Domain (RBD219-N1) Formulated with Alum Induces Protective Immunity and Reduces Immune Enhancement. bioRxiv. 10.1101/2020.05.15.098079

47. Zang, J., Gu, C., Zhou, B., Zhang, C., Yang, Y., Xu, S., Zhang, X., Zhou, Y., Bai, L., Wu, Y., Sun, Z., Zhang, R., Deng, Q., Yuan, Z., Tang, H., Qu, D., Lavillette, D., Xie, Y., and Huang, Z. (2020) Immunization with the receptor-binding domain of SARS-CoV-2 elicits antibodies cross-neutralizing SARS-CoV-2 and SARS-CoV without antibody-dependent enhancement. bioRxiv. 10.1101/2020.05.21.107565

48. Ren, A. W., Sun, H., Gao, G. F., Chen, J., Sun, S., Zhao, R., Gao, G., Hu, Y., Zhao, G., Chen, Y., Jin, X., Fang, F., Chen, J., Wang, Q., Gong, S., Gao, W., Sun, Y., Su, J., Sun, L., and Ph, D. (2020) Recombinant SARS-CoV-2 spike S1-Fc fusion protein induced high levels of neutralizing responses in nonhuman primates. bioRxiv. 10.1101/2020.04.21.052209

49. Dai, L., Zheng, T., Xu, K., Han, Y., Xu, L., Huang, E., An, Y., Cheng, Y., Li, S., Liu, M., Yang, M., Li, Y., Cheng, H., Yuan, Y., Zhang, W., Ke, C., Wong, G., Qi, J., Qin, C., Yan, J., and Gao, G. F. (2020) A Universal Design of Betacoronavirus Vaccines against COVID-19, MERS, and SARS. Cell. 182, 722–733.e11

50. Quinlan, B. D., Mou, H., Zhang, L., Guo, Y., He, W., Ojha, A., Parcells, M. S., Luo, G., Li, W., Zhong, G., Choe, H., and Farzan, M. (2020) The SARS-CoV-2 receptor-binding domain elicits a potent neutralizing response without antibody-dependent enhancement. bioRxiv. 10.1101/2020.04.10.036418

51. Powell, A. E., Zhang, K., Sanyal, M., Tang, S., Weidenbacher, P. A., Li, S., Pham, T. D., Pak, J. E., Chiu, W., and Kim, P. S. (2020) A single immunization with spike-functionalized ferritin vaccines elicits neutralizing antibody responses against SARS-CoV-2 in mice. bioRxiv. 10.1101/2020.08.28.272518

52. Walls, A. C., Fiala, B., Schäfer, A., Wrenn, S., Pham, M. N., Murphy, M., Tse, L. V, Shehata, L., O’Connor, M. A., Chen, C., Navarro, M. J., Miranda, M. C., Pettie, D., Ravichandran, R., Kraft, J. C., Ogohara, C., Palser, A., Chalk, S., Lee, E.-C., Kepl, E., Chow, C. M., Sydeman, C., Hodge, E. A., Brown, B., Fuller, J. T., Dinnon, K. H., Gralinski, L. E., Leist, S. R., Gully, K. L., Lewis, T. B., Guttman, M., Chu, H. Y., Lee, K. K., Fuller, D. H., Baric, R. S., Kellam, P., Carter, L., Pepper, M., Sheahan, T. P., Veesler, D., and King, N. P. (2020) Elicitation of potent neutralizing antibody responses by designed protein nanoparticle vaccines for SARS-CoV-2. bioRxiv. 10.1101/2020.08.11.247395

53. Zhang, N.-N., Li, X.-F., Deng, Y.-Q., Zhao, H., Huang, Y.-J., Yang, G., Huang, W.-J., Gao, P., Zhou, C., Zhang, R.-R., Guo, Y., Sun, S.-H., Fan, H., Zu, S.-L., Chen, Q., He, Q., Cao, T.-S., Huang, X.-Y., Qiu, H.-Y., Nie, J.-H., Jiang, Y., Yan, H.-Y., Ye, Q., Zhong, X., Xue, X.-L., Zha, Z.-Y., Zhou, D., Yang, X., Wang, Y.-C., Ying, B., and Qin, C.-F. (2020) A Thermostable mRNA Vaccine against COVID-19. Cell. 182, 1271–1283.e16

54. Yang, J., Wang, W., Chen, Z., Lu, S., Yang, F., Bi, Z., Bao, L., Mo, F., Li, X., Huang, Y., Hong, W., Yang, Y., Zhao, Y., Ye, F., Lin, S., Deng, W., Chen, H., Lei, H., Zhang, Z., Luo, M., Gao, H., Zheng, Y., Gong, Y., Jiang, X., Xu, Y., Lv, Q., Li, D., Wang, M., Li, F., Wang, S., Wang, G., Yu, P., Qu, Y., Yang, L., Deng, H., Tong, A., Li, J., Wang, Z., Yang, J., Shen, G., Zhao, Z., Li, Y., Luo, J., Liu, H., Yu, W., Yang, M., Xu, J., Wang, J., Li, H., Wang, H., Kuang, D., Lin, P., Hu, Z., Guo, W., Cheng, W., He, Y., Song, X., Chen, C., Xue, Z., Yao, S., Chen, L., Ma, X., Chen, S., Gou, M., Huang, W., Wang, Y., Fan, C., Tian, Z., Shi, M., Wang, F.-S., Dai, L., Wu, M., Li, G., Wang, G., Peng, Y., Qian, Z., Huang, C., Lau, J. Y.-N., Yang, Z., Wei, Y., Cen, X., Peng, X., Qin, C., Zhang, K., Lu, G., and Wei, X. (2020) A vaccine targeting the RBD of the S protein of SARS-CoV-2 induces protective immunity. Nature. 10.1038/s41586-020-2599-8

55. Kesavardhana, S., Das, R., Citron, M., Datta, R., Ecto, L., Srilatha, N. S., DiStefano, D., Swoyer, R., Joyce, J. G., Dutta, S., LaBranche, C. C., Montefiori, D. C., Flynn, J. A., and Varadarajan, R. (2017) Structure-based Design of Cyclically Permuted HIV-1 gp120 Trimers That Elicit Neutralizing Antibodies. J. Biol. Chem. 292, 278–291

56. Bell, J. M., Chen, M., Baldwin, P. R., and Ludtke, S. J. (2016) High resolution single particle refinement in EMAN2.1. Methods. 100, 25–34

57. Reboul, C. F., Eager, M., Elmlund, D., and Elmlund, H. (2018) Single-particle cryo-EM-Improved ab initio 3D reconstruction with SIMPLE/PRIME Protein Sci. 27, 51–61

58. Matsuura, Y., Tani, H., Suzuki, K., Kimura-Someya, T., Suzuki, R., Aizaki, H., Ishii, K., Moriishi, K., Robison, C. S., Whitt, M. A., and Miyamura, T. (2001) Characterization of Pseudotype VSV Possessing HCV Envelope Proteins. Virology. 286, 263–275

59. Joseph, J., T, K., Ajay, A., Das, V. R. A., and Raj, V. S. (2020) Green tea and Spirulina extracts inhibit SARS, MERS, and SARS-2 spike pseudotyped virus entry in vitro. bioRxiv. 10.1101/2020.06.20.162701

